# A D-2-hydroxyglutarate biosensor based on specific transcriptional regulator DhdR

**DOI:** 10.1101/2021.02.18.430539

**Authors:** Dan Xiao, Wen Zhang, Xiaoting Guo, Yidong Liu, Chunxia Hu, Shiting Guo, Zhaoqi Kang, Xianzhi Xu, Cuiqing Ma, Chao Gao, Ping Xu

## Abstract

D-2-Hydroxyglutarate (D-2-HG) is a metabolite in many physiological metabolic processes. When D-2-HG is aberrantly accumulated due to mutations in isocitrate dehydrogenases or D-2-HG dehydrogenase, it functions in a pro-oncogenic manner and is thus considered a therapeutic target and biomarker in many cancers. In this study, DhdR from *Achromobacter denitrificans* NBRC 15125 was identified as an allosteric transcription factor that negatively regulates D-2-HG dehydrogenase expression and responds to presence of D-2-HG. It is the first known transcription regulator specifically responding to D-2-HG across all domains of life. Based on the allosteric effect of DhdR, a D-2-HG biosensor was developed by combining DhdR with amplified luminescent proximity homogeneous assay technology. The biosensor was able to detect D-2-HG in serum, urine, and cell culture with high specificity and sensitivity. Additionally, this biosensor was also successfully used to identify the role of D-2-HG metabolism in lipopolysaccharide biosynthesis of *Pseudomonas aeruginosa*, demonstrating its broad usages.

## Introduction

D-2-Hydroxyglutarate (D-2-HG) has traditionally been considered an abnormal metabolite associated with the neurometabolic disorder D-2-hydroxyglutaric aciduria^1^. However, D-2-HG accumulation, due to mutations of isocitrate dehydrogenase (IDH), has been observed in many tumor cells, and thus D-2-HG is also considered an oncometabolite^2–6^. Recent work has revealed that D-2-HG may also be an important metabolic intermediate involved in different core metabolic processes, including L-serine synthesis^7^, lysine degradation^8^, and 4-hydroxybutyrate catabolism^9^. D-2-HG dehydrogenase (D2HGDH) catalyzes the conversion of D-2-HG to 2-ketoglutarate (2-KG) and is the key enzyme involved in D-2-HG catabolism^7^. D-2-HG is harmful to cells and is present at rather low levels under physiological conditions, suggesting that organisms may have evolved specific mechanisms to recognize D-2-HG accumulation and enhance D2HGDH expression to catabolize D-2-HG^10–13^. However, the regulation of D2HGDH expression is not fully understood and how organisms respond to the presence of D-2-HG has not yet been elucidated.

D-2-HG levels are increased in patients with D-2-hydroxyglutaric aciduria (D-2-HGA) and IDH mutation-related cancers^1, 2^. As such, the detection of D-2-HG is relevant for the diagnosis and monitoring of these diseases^14–16^. Gas chromatography-tandem mass spectrometry (GC-MS/MS)^17, 18^ and liquid chromatography-tandem mass spectrometry (LC-MS/MS)^19, 20^ are often used to quantitatively assess D-2-HG levels. However, chiral derivatization with proper reagents is necessary to distinguish D-2-HG from its mirror-image enantiomer L-2-HG, and these methods are time-consuming and laborious. Allosteric transcription factors (aTFs) in bacteria have evolved to sense a variety of chemicals^21^.

Various aTFs, such as HucR, HosA, TetR, and SRTF1, have been well characterized, and then coupled with different transduction systems to develop convenient biosensors to assay for uric acid, 4-hydroxybenzoic acid, tetracycline, and progesterone^22–25^. Until now, no aTF that specifically responds to D-2-HG has been identified, restricting the development of D-2-HG biosensors.

In this study, the D-2-HG catabolism regulator DhdR was identified in *Achromobacter denitrificans* NBRC 15125. This aTF can depress the expression of the D2HGDH-encoding gene *d2hgdh* and specifically responds to D-2-HG. Then, DhdR was combined with amplified luminescent proximity homogeneous assay (AlphaScreen) technology, a bead-based immunoassay, to develop a D-2-HG biosensor with high specificity and sensitivity. Various biological samples were selected to demonstrate the utility of the biosensor. In addition, the biosensor was used to identify the UDP-2-acetamido-2-deoxy-D-glucuronic acid (UDP-GlcNAcA) 3-dehydrogenase WbpB of *Pseudomonas aeruginosa* PAO1 as a D-2-HG anabolic protein that also participates in the intracellular generation of D-2-HG.

## Results

### Genome analysis predicts a D-2-HG catabolism operon

Bacteria have evolved various aTFs to respond to different stimuli. To identify the possible aTFs that respond to D-2-HG and regulate the expression of D2HGDH, two open reading frames (ORFs) upstream and two ORFs downstream of D2HGDH in bacteria containing the D2HGDH were subjected to gene occurrence profile analysis. GntR family transcriptional regulator, electron transfer flavoprotein (ETF), 3-phosphoglycerate dehydrogenase (SerA), carbohydrate diacid regulator (CdaR), lactate permease (LldP) and glycolate oxidase iron-sulfur subunit (GlcF) appear to be the most frequently observed in the neighborhood of D2HGDH (Supplementary Table 1).

Subsequent chromosomal gene clustering analysis identified five typical patterns of organized chromosomal clusters (Fig. 1a). GntR and CdaR are located upstream of D2HGDH in *Parageobacillus thermoglucosidasius* DSM 2542 and *Bacillus cereus* NJ-W, respectively. However, LldP and GlcF, responding for the metabolism of two other hydroxycarboxylic acids, lactate and glycolate, are also located adjacent to D2HGDH in *P. thermoglucosidasius* DSM 2542 and *B. cereus* NJ-W. Despite that these two transcriptional regulators might respond to D-2-HG and regulate D2HGDH expression, their possible effector non-specificities should not be overlooked. Interestingly, in *A. denitrificans* NBRC 15125, GntR is located directly upstream of D2HGDH and no adjacent hydroxycarboxylic acid metabolism-related protein is encoded. Thus, in *A. denitrificans* NBRC 15125, GntR and D2HGDH might comprise an operon that is specifically responsible for D-2-HG catabolism and warrant further investigation. The uncharacterized transcriptional regulator GntR was tentatively designated as **<u>D</u>**-2-**<u>h</u>**ydroxyglutarate **<u>d</u>**ehydrogenase **<u>r</u>**egulator, DhdR.

**Figure 1.**
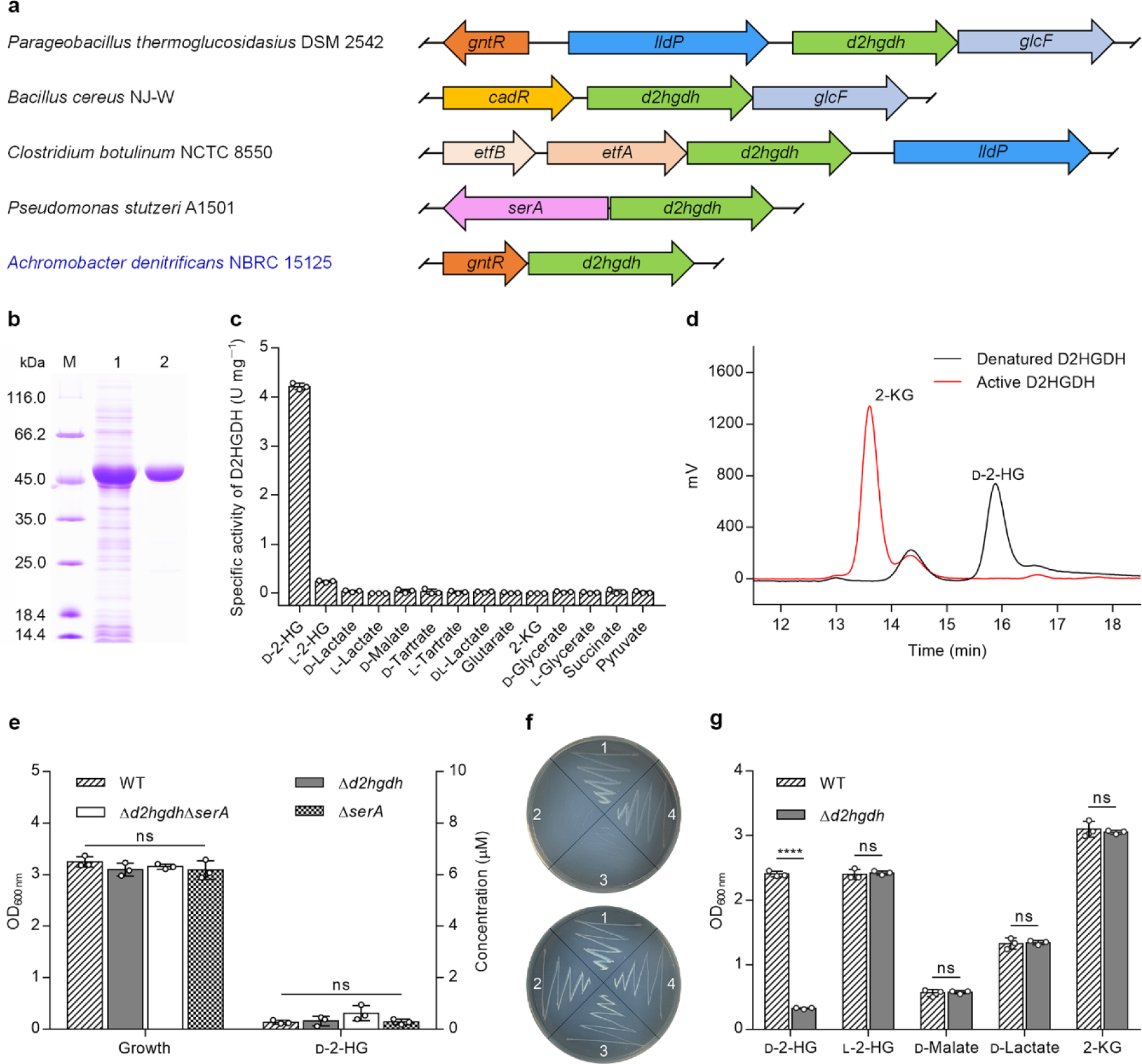
D2HGDH contributes to D-2-HG catabolism in *A. denitrificans* NBRC 15125. **a**, Schematic representation of genome context analysis of *d2hgdh* in different species. Orthologs are shown in the same color. Arrows indicate the direction of gene translation. The corresponding proteins encoded by the indicated genes are D-2-HG dehydrogenase (*d2hgdh*), GntR family regulator protein (*gntR*), lactate permease (*lldP*), glycolate oxidase iron-sulfur subunit (*glcF*), carbohydrate diacid regulator (*cadR*), electron transfer flavoprotein alpha/beta-subunit (*etfA/B*), 3-phosphoglycerate dehydrogenase (*serA*). **b**, SDS-PAGE analysis of purified D2HGDH in *A. denitrificans* NBRC 15125. Lane M, molecular weight markers; lane 1, crude extract of *E. coli* BL21(DE3) harboring pETDuet-*d2hgdh*; lane 2, purified D2HGDH using a HisTrap column. **c**, Substrate specificity of D2HGDH in *A. denitrificans* NBRC 15125. Data shown are mean ± standard deviations (s.d.) (*n* = 3 independent experiments). **d**, HPLC analysis of the product of D2HGDH-catalyzed D-2-HG dehydrogenation. The reaction mixtures containing D-2-HG (1 mM), 3-(4,5-dimethylthiazol-2-yl)-2,5-diphenyltetrazolium bromide (MTT, 5 mM) and active or denatured D2HGDH (1 mg mL^−1^) in 50 mM Tris-HCl (pH 7.4) were incubated at 37 °C for 5 min. Black line, the reaction with denatured D2HGDH; red line, the reaction with active D2HGDH; **e**, Growth and extracellular D-2-HG concentrations of *A. denitrificans* NBRC 15125 and its derivatives cultured in LB medium at mid-log stage. Data shown are mean ± s.d. (*n =* 3 independent experiments). **f**, Growth of *A. denitrificans* NBRC 15125 and its derivatives on solid minimal medium containing 5 g L^−1^ D-2-HG (top) or 2-KG (bottom) as the sole carbon source after 36 h. 1. *A. denitrificans* NBRC 15125. 2. *A. denitrificans* NBRC 15125 (Δ*d2hgdh*); 3. *A. denitrificans* NBRC 15125 (Δ*d2hgdh*Δ*serA*); 4. *A. denitrificans* NBRC 15125 (Δ*serA*). **g**, Growth of *A. denitrificans* NBRC 15125 and *A. denitrificans* NBRC 15125 (Δ*d2hgdh*) in minimal medium containing different carbon sources after 24 h. Data shown are mean ± s.d. (*n* = 3 independent experiments). The significance was analyzed by a two-tailed, unpaired *t* test. ****, *P* < 0.0001; ns, no significant difference (*P* ≥ 0.05).

### D2HGDH is required for extracellular D-2-HG metabolism

D2HGDH of *A. denitrificans* NBRC 15125 was expressed as a His-tagged protein in *Escherichia coli* BL21(DE3) and purified by affinity chromatography. Sodium dodecyl sulfate-polyacrylamide gel electrophoresis (SDS-PAGE) and size exclusion chromatography revealed that D2HGDH was a dimer (Fig. 1b and Supplementary Fig. 1). Substrate screening revealed that D2HGDH had narrow substrate specificity and only exhibited distinct activity towards D-2-HG (Fig. 1c). The product of D2HGDH-catalyzed dehydrogenation of D-2-HG was identified to be 2-KG by high performance liquid chromatography (HPLC) analysis (Fig. 1d). The apparent *K*_m_ and *V*_max_ of purified D2HGDH for D-2-HG were 31.16 ± 1.40 μM and 3.95 ± 0.09 U mg^−1^, respectively (Supplementary Fig. 2).

SerA is the key enzyme for L-serine biosynthesis in various species, such as *P. stutzeri* A1501, *P. aeruginosa*^7^, *E. coli*^26^, *Saccharomyces cerevisiae*^27^, and *Homo sapiens*^28^, and can catalyze the conversion of 2-KG to D-2-HG. SerA of *A. denitrificans* NBRC 15125 was over-expressed, purified, and identified to be a dimer (Supplementary Fig. 3a,b). It was also able to reduce 2-KG to D-2-HG (Supplementary Fig. 3c,d), and the apparent *K*_m_ and *V*_max_ for 2-KG were 57.03 ± 2.12 μM and 1.96 ± 0.03 U mg^−1^, respectively (Supplementary Fig. 3e). However, *A. denitrificans* NBRC 15125 (Δ*d2hgdh*), *A. denitrificans* NBRC 15125 (Δ*serA*), and *A. denitrificans* NBRC 15125 (Δ*d2hgdh*Δ*serA*) exhibited no growth defects in Luria-Bertani (LB) medium and no accumulation of D-2-HG was detected (Fig. 1e and Supplementary Fig. 4). These results demonstrated that there is no SerA-catalyzed intracellular D-2-HG production in *A. denitrificans* NBRC 15125 and D2HGDH does not participate in intracellular D-2-HG metabolism. *A. denitrificans* NBRC 15125 (Δ*serA*) and *A. denitrificans* NBRC 15125 (Δ*d2hgdh*Δ*serA*) are able to grow well on solid minimal medium without exogenous L-serine (Fig. 1f). It should be noted that *A. denitrificans* NBRC 15125 is able to use D-2-HG as the sole carbon source for growth. Disruption of *d2hgdh* abolished the ability of *A. denitrificans* NBRC 15125 (Δ*d2hgdh*) to assimilate D-2-HG but did not affect growth in the presence of L-2-HG, D-malate, D-lactate, and 2-KG, indicating that D2HGDH is critical for extracellular D-2-HG utilization in this strain (Fig. 1g).

### DhdR represses *d2hgdh* expression and responses to D-2-HG

The *dhdR* and *d2hgdh* genes are adjacent to each other in the genome of *A. denitrificans* NBRC 15125. The intergenic region between *dhdR* and *d2hgdh* was amplified by reverse transcription-PCR (RT-PCR) using cDNA generated from the total RNA of cells grown in the presence of D-2-HG (Fig. 2a), suggesting that these two genes are co-transcribed. The transcription initiation site of *dhdR* was identified by rapid amplification of cDNA ends. The transcription initiation site was confirmed to be an adenine (A) residue that overlapped with the *dhdR* start codon, with the putative −10 (TATAAT) and −35 (TTATCA) regions being separated by 20 bp (Fig. 2b).

**Figure 2.**
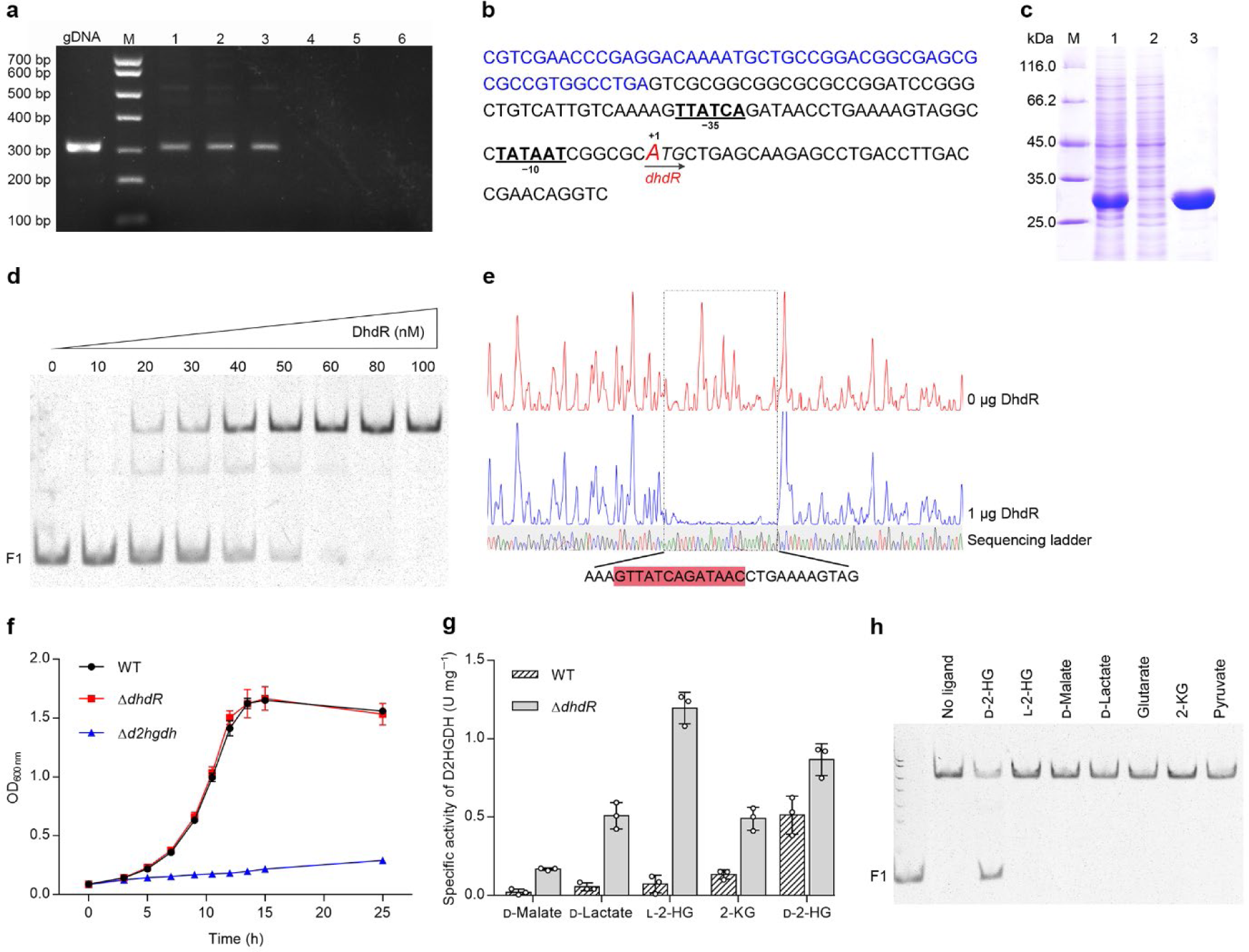
DhdR negatively regulates the catabolism of D-2-HG. **a**, Identification of the cotranscription of *dhdR* and *d2hgdh* by RT-PCR. Genomic DNA of *A. denitrificans* NBRC 15125 was used as a positive control (left lane). RT-PCR was performed by using the cDNA as template (lane 1-3). RNA was used as negative control (lane 4-6). **b**, Map of the intergenic region upstream of *dhdR*-*d2hgdh* operon. Transcriptional initiation site is shown in enlarged red letter and marked by +1. The putative −35 and −10 regions are shown in bold and underlined. The translational start codon of *dhdR* is shown in italics. **c**, SDS-PAGE analysis of purified DhdR. Lane M, molecular weight markers; lane 1, crude extract of *E. coli* BL21(DE3) harboring pETDuet-*dhdR*; lane 2, the unbound protein of the HisTrap HP column; lane 3, purified DhdR using a HisTrap column. **d**, EMSAs with the 81-bp fragment (F1) upstream of *dhdR* (10 nM) and purified DhdR (0, 10, 20, 30, 40, 50, 60, 80, and 100 nM). **e**, DNase I footprinting analysis of DhdR binding to the *dhdR* promoter region. F1 was labeled with FAM dye and incubated with 1 μg DhdR (blue line) or without DhdR (red line). Sequences of DhdR protected region are shown at the bottom and the palindrome sequences are indicated with a red box. **f**, Growth of *A. denitrificans* NBRC 15125 and its derivatives cultured in minimal medium containing 3 g L^−1^ D-2-HG. Data shown are mean ± s.d. (*n =* 3 independent experiments). **g**, Activity of D2HGDH in *A. denitrificans* NBRC 15125 and its derivatives cultured in minimal medium with different compounds as the sole carbon source. Data shown are mean ± s.d. (*n =* 3 independent experiments). **h**, D-2-HG prevents DhdR binding to F1. EMSAs were performed with F1 (10 nM) and a 7-fold molar excess of DhdR dimer in the absence or presence of 40 mM D-2-HG, L-2-HG, D-malate, D-lactate, glutarate, 2-KG, or pyruvate. The leftmost lane shows the migration of free DNA.

The His_6_-tagged DhdR was expressed and purified by affinity chromatography (Fig 2c). Size exclusion chromatography and SDS-PAGE indicated that DhdR behaves as a dimer (Supplementary Fig. 5). An electrophoretic mobility shift assay (EMSA) was performed using purified DhdR and the 81-bp fragment (F1) upstream of *dhdR*. As shown in Fig. 2d, DhdR formed a complex with F1 in a concentration-dependent manner and a complete gel shift was observed with 6-fold molar excess of DhdR. A DNase I footprinting assay was also performed using purified DhdR and F1 end-labeled with 6-carboxyfluorescein (FAM) and one clearly protected region containing a motif with a palindromic sequence (5′-GTTATCAGATAAC-3′) was identified (Fig. 2e). The protected region overlapped the putative −35 region relative to the transcription initiation site of *dhdR*, implying that DhdR functions as a repressor.

As shown in Fig. 2f, *A. denitrificans* NBRC 15125 (Δ*dhdR*) was able to grow in the presence of D-2-HG, as was the wild-type strain, whereas *A. denitrificans* NBRC 15125 (Δ*d2hgdh*) was not. Additionally, the D2HGDH activity of *A. denitrificans* NBRC 15125 (Δ*dhdR*) cultured in medium containing D-malate, D-lactate, L-2-HG, or 2-KG was much higher than that of *A. denitrificans* NBRC 15125, further supporting that DhdR represses D2HGDH expression (Fig. 2g).

D2HGDH activity in *A. denitrificans* NBRC 15125 was significantly increased when cultured in medium containing D-2-HG (Fig. 2g), suggesting that D-2-HG can induce D2HGDH expression. The effects of L-2-HG, D-malate, D-lactate, glutarate, 2-KG, pyruvate, and D-2-HG on the binding of DhdR to F1 were also assessed by EMSA. As shown in Fig. 2h, only D-2-HG could prevent the binding of DhdR to F1. Therefore, DhdR negatively regulates D-2-HG catabolism and D-2-HG is the specific effector of DhdR.

### DhdR behaves as a recognition element to develop biosensors

The specific response of DhdR to D-2-HG inspired us to develop a D-2-HG biosensor with DhdR as the biorecognition element (Fig. 3a). The AlphaScreen technology was selected as the transducing element of this biosensor. As shown in Fig. 3b, a biotinylated DNA fragment, Bio-*dhdO*, containing the DhdR protected region (5’-GTTATCAGATAAC-3’) and His_6_-tagged DhdR are bound to streptavidin donor and nickel chelate acceptor beads, respectively. D-2-HG can prevent the binding of DhdR to Bio-*dhdO* and increase the distance from donor beads to acceptor beads. Therefore, the singlet oxygen (^1^O2) generated from the conversion of ambient oxygen upon laser excitation of donor beads at 680 nm cannot be transferred to acceptor beads, resulting in reduced luminescence signals at 520–620 nm. Through this approach, the concentration of D-2-HG can be connected with the measurable luminescence signals. A formal D-2-HG measurement process includes diluting the biological samples (if necessary), incubating Bio-*dhdO* and His_6_-tagged DhdR with commercial streptavidin donor and nickel chelate acceptor beads, and detecting luminescence signals with a laboratory plate reader (Supplementary Fig. 6).

**Figure 3.**
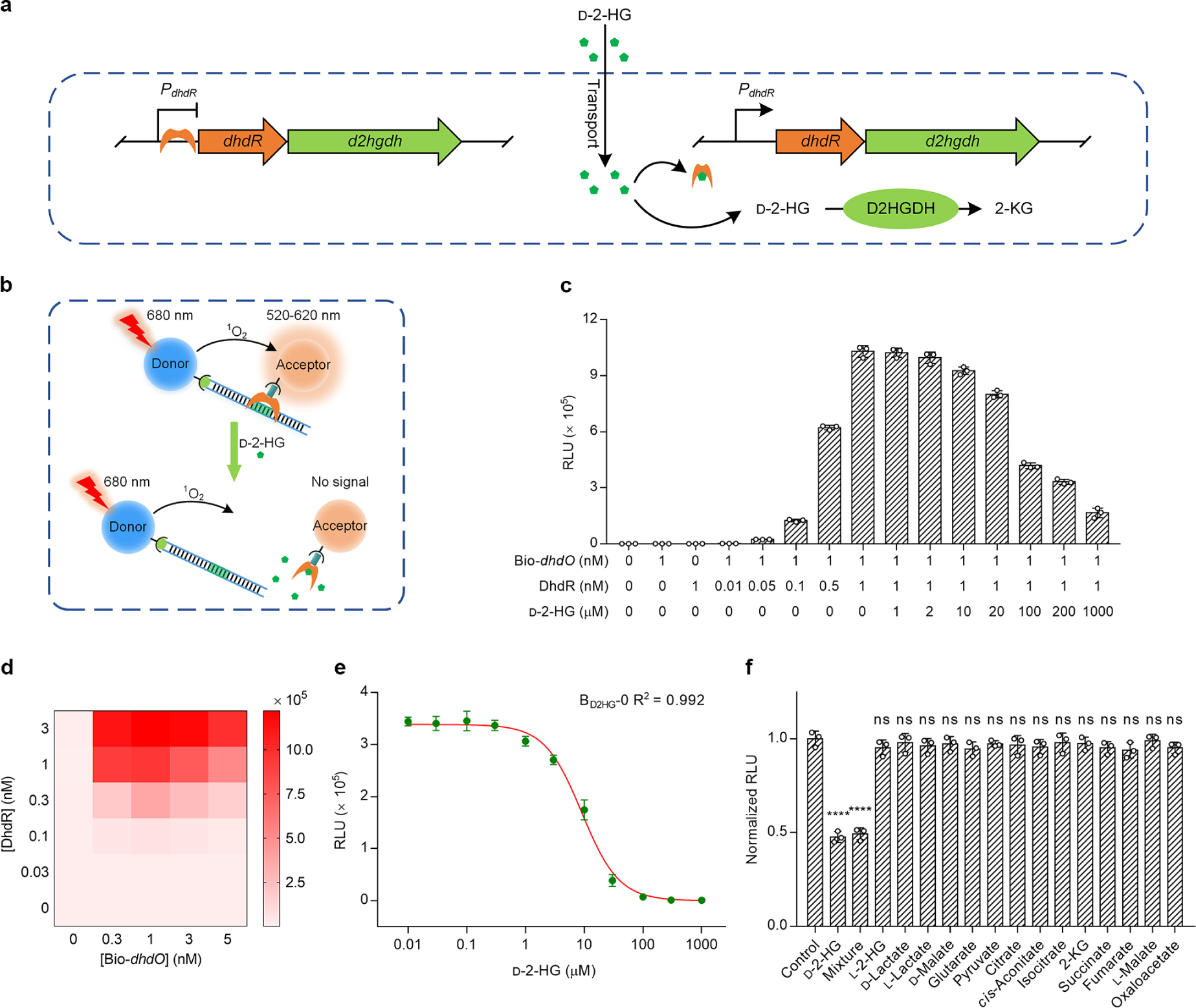
Design of the D-2-HG biosensor (B_D2HG_). **a**, Proposed model for the regulation of D-2-HG catabolism by DhdR in *A. denitrificans* NBRC 15125. DhdR represses the expression of *dhdR-d2hgdh* genes. D-2-HG is the effector of DhdR and prevents DhdR binding to the *dhdR* promoter region. **b**, Schematic diagram of the biosensor based on DhdR and AlphaScreen. **c**, Evaluation of allosteric effect of D-2-HG on *dhdO*-DhdR interaction by AlphaScreen. Data shown are mean ± s.d. (*n =* 3 independent experiments). **d**, Determination of the optimal concentrations of DhdR and Bio-*dhdO* by cross-titration. Each heat map cell shows the average of the luminescence signals of each titration (*n =* 3 independent experiments). **e**, Response of B_D2HG_-0 to different concentrations of D-2-HG. **f**, Specificity of B_D2HG_-0. The concentrations of tested chemicals were 10 μM. The column mixture contained 10 μM D-2-HG and 10 μM all other tested chemicals. The corresponding luminescence signals are normalized to the maximal value (luminescence signals of control) in the absence of D-2-HG. Data shown are mean ± s.d. (*n =* 3 independent experiments). The significance was analyzed by a two-tailed, unpaired *t* test. ****, *P* < 0.0001; ns, no significant difference (*P* ≥ 0.05).

To demonstrate the feasibility of this approach, the oscillations of relative luminescence unit (RLU) induced by *dhdO*-DhdR interaction were assayed. As shown in Fig. 3c, almost no luminescence signal was observed in the reaction system with only 1 nM Bio-*dhdO*. Upon the addition of His_6_-tagged DhdR, the luminescence signal appeared and increased as the concentration of His_6_-tagged DhdR increased (0.01–1 nM). The addition of extra D-2-HG to induce the dissociation of Bio-*dhdO* and His_6_-tagged DhdR led to a significantly decreased luminescence intensity, as expected. Then, the reaction system for the quantification of D-2-HG was optimized. HBS-P buffer, which gives the highest signal to noise ratio (S/N) and lowest background signal (Supplementary Fig. 7), was determined to be the optimal working buffer for D-2-HG quantification. Additionally, this buffer also has a high buffering capacity across a range sample pH values (pH 4.0–10.0) and various inorganic substances or amino acids (100 μM) did not interfere with the RLU of the biosensor (Supplementary Fig. 8 and Supplementary Table 2). Next, the concentrations of DhdR and Bio-*dhdO* were optimized through cross-titration. As shown in Fig. 3d, a hook effect appeared when the concentration of Bio-*dhdO* was higher than 1 nM. A credible signal (3.7 × 10^5^ RLU, S/N = 151.2) could be acquired when the concentrations of DhdR and Bio-*dhdO* were 0.3 nM and 1 nM, respectively. Thus, 0.3 nM of DhdR and 1 nM of Bio-*dhdO* were used in the original D-2-HG biosensor B_D2HG_-0 and the luminescence signals emitted at a series of D-2-HG concentrations were measured. As shown in Fig. 3e, B_D2HG_-0 responded to D-2-HG supplementation in a dose-dependent manner and the limit of detection (LOD) of B_D2HG_-0 was calculated to be 0.80 μM. B_D2HG_-0 exhibited good specificity and did not respond to D-2-HG analogues (L-2-HG, D-lactate, L-lactate, D-malate, and glutarate) or metabolites involved in the tricarboxylic acid cycle (pyruvate, citrate, *cis*-aconitate, isocitrate, 2-KG, succinate, fumarate, L-malate, and oxaloacetate). In addition, a mixture of these chemicals also failed to interfere with the response of B_D2HG_-0 to D-2-HG (Fig. 3f).

### Mutation of binding site results in a sensitive biosensor

Reducing the affinity between DhdR and its binding site could increase the equilibrium dissociation constant *K*_D_ and finally improve the sensitivity and linear detection range of the biosensor. Thirteen mutations were generated through displacing one or two bases in the hyphenated dyad symmetry of the DhdR binding site (DBS). Isothermal titration calorimetry (ITC) was used to assess the *K*_D_ values of DhdR for the thirteen mutated DBSs (Supplementary Fig. 9). As shown in Fig. 4a, the *K*_D_ values of DhdR for the DBS mutants D1, D2, D8, and D12 were higher than those of other mutants (Fig. 4a). These four mutants were then used to construct the biosensors B_D2HG_-1, B_D2HG_-2, B_D2HG_-8, and B_D2HG_-12.

**Figure 4.**
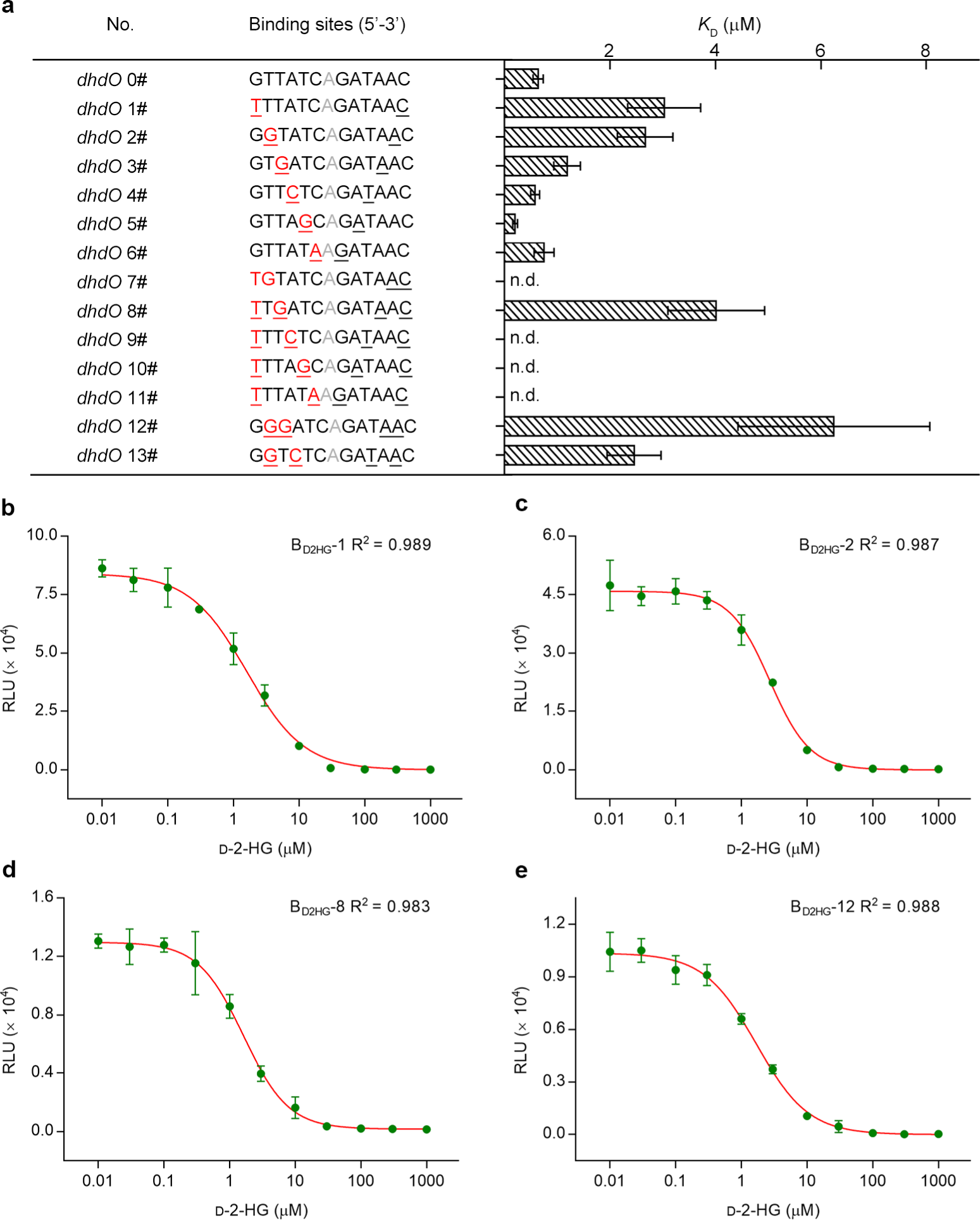
Optimization of the D-2-HG biosensor (B_D2HG_). **a**, Influence of point mutations in DBS on affinity determined by ITC. The point mutation is shown in red letter. The 1-bp interval in palindrome sequence is shown in gray letter. n.d., not detected. **b-e**, Responses of the optimized biosensors to different concentrations of D-2-HG. **b**, Dose-response curve of B_D2HG_-1; **c**, Dose-response curve of B_D2HG_-2; **d**, Dose-response curve of B_D2HG_-8; **e**, Dose-response curve of B_D2HG_-12. Data shown are mean ± s.d. (*n =* 3 independent experiments).

As shown in Fig 4b-e, the four biosensors exhibited different responses to D-2-HG. Increased *K*_D_ values of DhdR for the DBS through point mutation could decrease the *I*_50_ (concentration of D-2-HG producing a 50% signal reduction) of biosensors (Supplementary Table 3). Among the four optimized biosensors, B_D2HG_-1 exhibited the lowest LOD (0.08 μM) and a wide linear range (0.3–20 μM) (Fig 4b-e and Supplementary Fig. 10 and Supplementary Table 3). This biosensor was selected to quantify D-2-HG levels in subsequent experiments.

### Performance of B_D2HG_-1 in D-2-HG quantitation

The concentration of D-2-HG in body fluids has the potential to be a clinical indicator of D-2-HG related diseases^1, 3, 29^. Considering the high sensitivity of B_D2HG_-1 for D-2-HG, small amounts of body fluids are required for D-2-HG quantitation and the dose-response curves of D-2-HG were determined in 100-fold diluted human serum or urine (Fig. 5a,b). Then, D-2-HG, at a range of concentrations, was spiked into human serum or urine and quantified using B_D2HG_-1 and LC-MS/MS. The agreement between results from LC-MS/MS and the D-2-HG biosensor were satisfactory (Fig. 5c,d). 2-KG, glutamate, glutamine, and isocitrate are possible precursors of D-2-HG *in vivo*. These metabolites and L-2-HG, the enantiomer of D-2-HG, do not decrease the luminescence signals of the biosensor. The addition of 2-KG, glutamate, glutamine, isocitrate, or L-2-HG in the assay system does not interfere D-2-HG quantitation in serum or urine (Fig. 5e,f).

**Figure 5.**
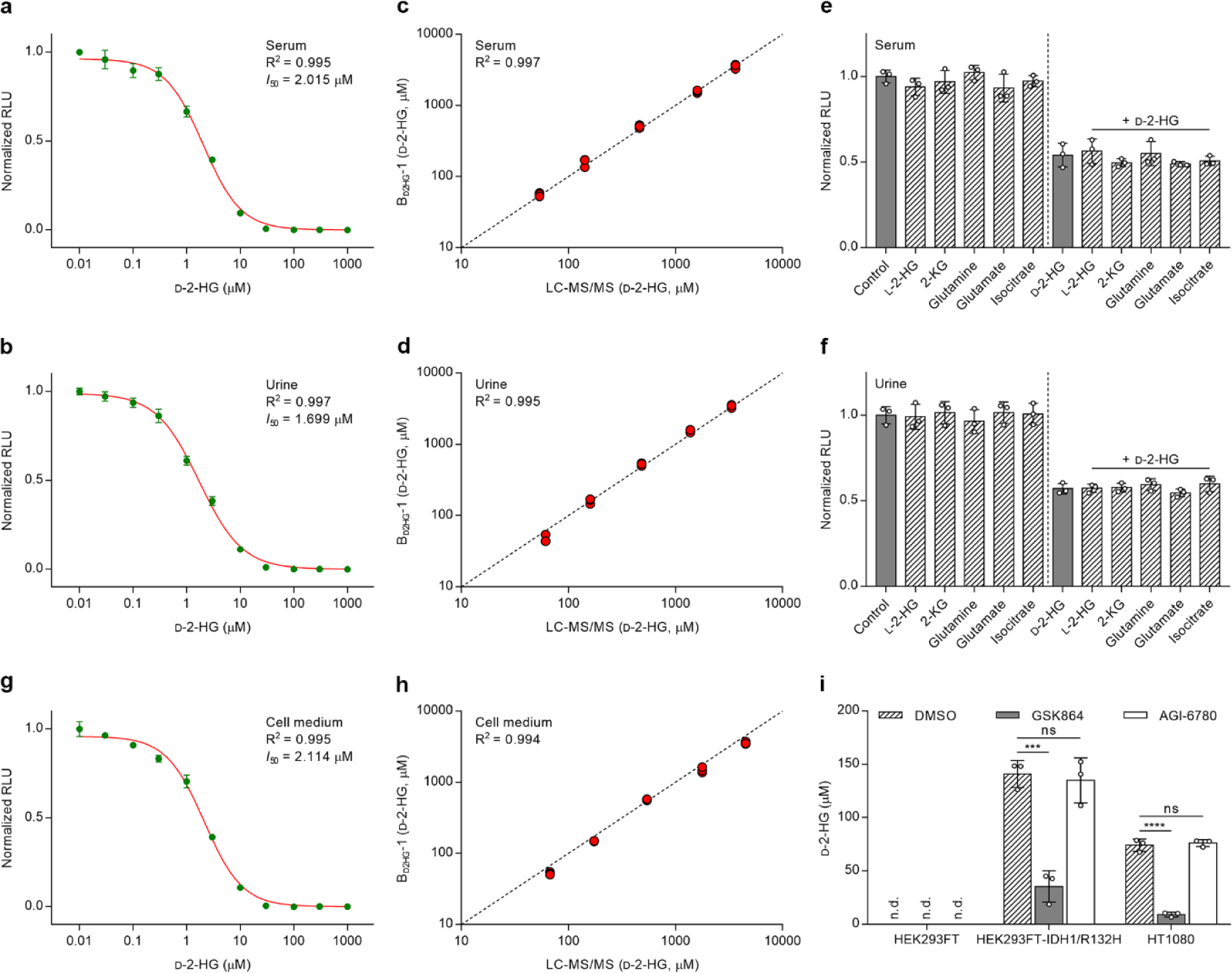
Measuring D-2-HG concentrations in various biological samples by using B_D2HG_-1. **a**, Normalized dose-response curve of B_D2HG_-1 in 100-fold diluted human serum. **b**, Normalized dose-response curve of B_D2HG_-1 in 100-fold diluted urine. **c**, Comparisons of analysis of D-2-HG in human serum by LC-MS/MS and B_D2HG_-1. **d**, Comparisons of analysis of D-2-HG in urine by LC-MS/MS and B_D2HG_-1. Black dotted line indicates a reference line. **e**, Effect of possible precursors and enantiomer of D-2-HG on the detection of D-2-HG in serum by B_D2HG_-1. **f**, Effect of possible precursors and enantiomer of D-2-HG on the detection of D-2-HG in urine by B_D2HG_-1. **e**,**f**, Metabolites were added in the assay systems with or without D-2-HG at physiologic concentrations (D-2-HG 1 mM, L-2-HG 1 mM, 2-KG 100 μM, glutamine 4 mM, glutamate 4 mM, and isocitrate 50 μM). **g**, Normalized dose-response curve of B_D2HG_-1 in 10-fold diluted cell culture medium. **h**, Comparisons of analysis of D-2-HG in cell medium by LC-MS/MS and B_D2HG_-1. Black dotted line indicates a reference line. **i**, D-2-HG detection in cell culture medium of different cells upon treatment with 0.5 μM GSK 864 or AGI-6780. DMSO was added as the control. The corresponding luminescence signals are normalized to the maximal value in the absence of D-2-HG. n.d., not detected. Data shown are mean ± s.d. (*n =* 3 independent experiments). The significance was analyzed by a two-tailed, unpaired *t* test. ***, *P* < 0.001; ****, *P* < 0.0001; ns, no significant difference (*P* ≥ 0.05).

The D-2-HG biosensor was also applied to quantify the generation of D-2-HG in different cell lines. The dose-response curve for D-2-HG was determined in 10-fold diluted cell culture medium and the biosensor B_D2HG_-1 also produced highly accurate quantification of D-2-HG (Fig. 5g,h). As shown in Fig. 5i, the D-2-HG biosensor B_D2HG_-1 could detect the increase in the levels of D-2-HG in the cell culture medium during the growth of HEK293FT cells transfected with IDH1/R132H and of HT1080 cells, a fibrosarcoma cell line carrying an IDH1/R132C mutation^30^. No D-2-HG was detected in the cell culture medium of HEK293FT cells. GSK864 and AGI-6780 are small molecules that selectively inhibit the cancer-associated mutants of IDH1 and IDH2/R140Q, respectively^31, 32^. As expected, treatment with 0.5 μM GSK864 significantly reduced the amount of extracellular D-2-HG of HEK293FT-IDH1/R132H cells and HT1080 cells, while AGI-6780 produced no detectable effect (Fig. 5i).

### D-2-HG metabolism in LPS synthesis in *P. aeruginosa* PAO1

Finally, the D-2-HG biosensor was used to investigate D-2-HG metabolism in *P. aeruginosa* PAO1, an important opportunistic pathogen that causes serious infections^33^. Based on the dose-response curves of D-2-HG in bacterial minimal medium, the growth and D-2-HG production of *P. aeruginosa* PAO1 and its derivative were assayed (Fig. 6a and Supplementary Figs. 11 and 12). In *P. aeruginosa* PAO1, SerA and D2HGDH (PA0317) were previously considered to be the key enzymes responsible for D-2-HG anabolism and catabolism, respectively. Deletion of *PA0317* caused extracellular D-2-HG accumulation in *P. aeruginosa* PAO1 (Δ*PA0317*)^7^. However, quantification of D-2-HG with the biosensor indicated that *P. aeruginosa* PAO1 (Δ*PA0317*Δ*serA*) still accumulated D-2-HG to a certain extent.

**Figure 6.**
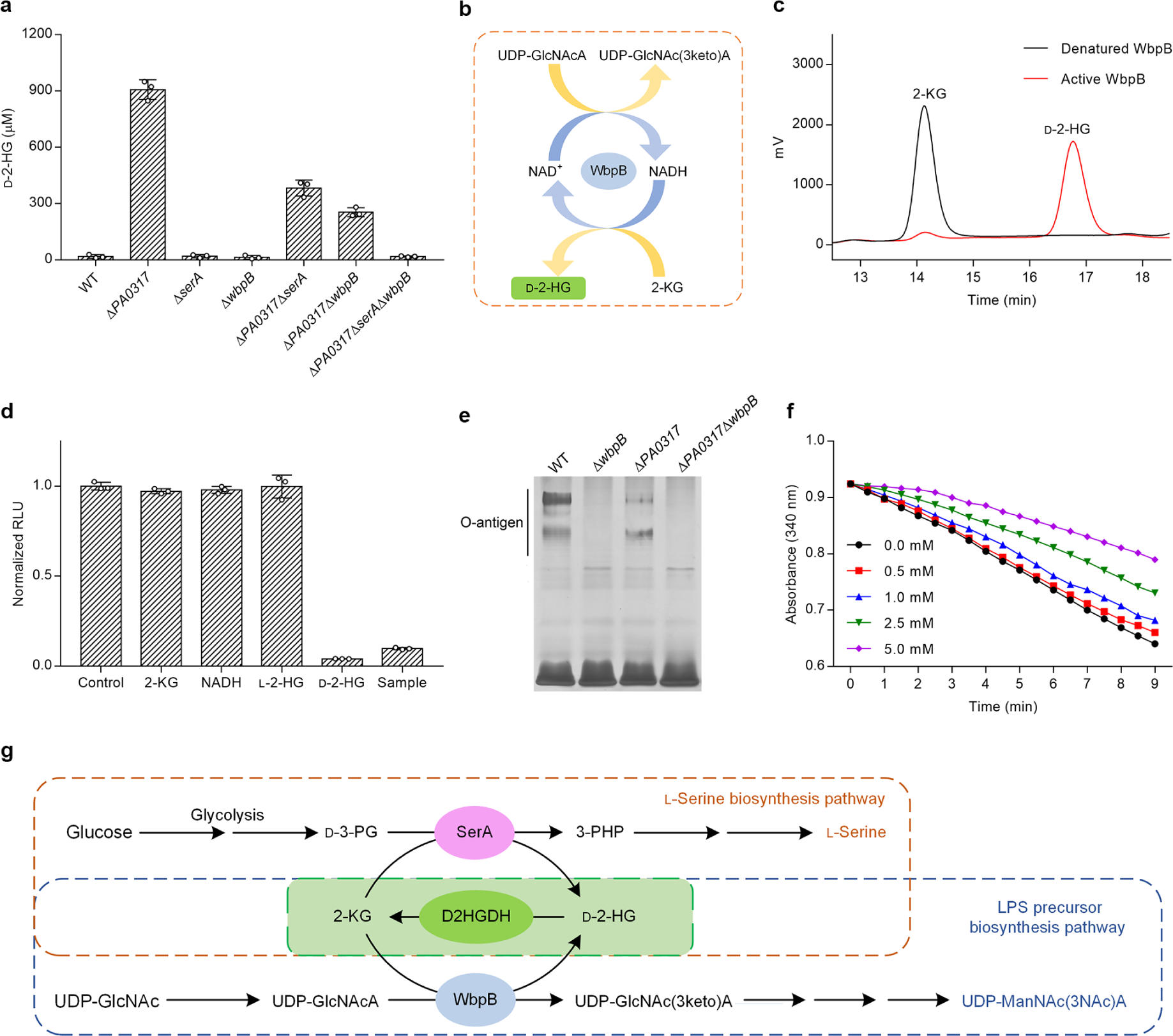
Identification of the D-2-HG metabolism pathways in *P. aeruginosa* PAO1 using B_D2HG-1_. **a**, Determination of extracellular D-2-HG of *P. aeruginosa* PAO1 and its derivatives by B_D2HG_-1. Data shown are mean ± s.d. (*n =* 3 independent experiments). **b**, Schematic diagram of WbpB-catalyzed coupled reaction. **c**, HPLC analysis of the product of WbpB-catalyzed 2-KG reduction. The reaction mixtures containing 2-KG (5 mM), NADH (5 mM) and active or denatured WbpB (0.4 mg mL^−1^) in 50 mM Tris-HCl (100 mM NaCl, pH 7.4) were incubated at 37 °C for 3 h. Black line, the reaction with denatured WbpB; red line, the reaction with active WbpB. **d**, Chiral analysis of WbpB produced 2-HG by B_D2HG_-1. The concentration of 2-KG, NADH, L-2-HG and D-2-HG is 20 μM. Sample is 50-fold diluted WbpB-catalyzed product. **e**, SDS-PAGE analysis of LPS of *P. aeruginosa* PAO1 and its derivatives. **f**, Inhibitory of D-2-HG toward WbpB. The reaction mixtures containing WbpB (0.3 mg mL^−1^), 2-KG (0.5 mM), NADH (200 μM) and 0−5 mM D-2-HG 50 mM Tris-HCl (100 mM NaCl, pH 7.4) were incubated at 37 °C for 9 min. **g**, D-2-HG metabolism pathways in *P. aeruginosa* PAO1. The coupling between D2HGDH and SerA or D2HGDH and WbpB makes the robust interconversion between D-2-HG and 2-KG, and facilitates the biosynthesis of L-serine (orange dotted box) and precursor of LPS in *P. aeruginosa* PAO1 (blue dotted box).

Lipopolysaccharide (LPS) functions as a natural barrier against antibiotics and plays a key role in the host-pathogen interactions of *P. aeruginosa*^34^. The synthesis of LPS in *P. aeruginosa* PAO1 involves the dehydrogenation of UDP-GlcNAcA to UDP-2-acetamido-2-deoxy-D-*ribo*-hex-3-uluronic acid [UDP-GlcNAc(3keto)A], which is catalyzed by UDP-GlcNAcA 3-dehydrogenase WbpB (Fig. 6b). As shown in Fig. 6a, *P. aeruginosa* PAO1 (Δ*PA0317*Δ*serA*Δ*wbpB*) no longer accumulated D-2-HG, suggesting that WbpB might also be involved in D-2-HG anabolism in *P. aeruginosa* PAO1.

Then, WbpB of *P. aeruginosa* PAO1 was over-expressed and purified (Supplementary Fig. 13). The spectral signal peak at 334 nm of purified WbpB indicated that this enzyme binds NADH tightly (Supplementary Fig. 14). When 2-KG was added to a solution with purified WbpB, the tightly bound NADH was oxidized and absorbance at 334 nm dramatically decreased (Supplementary Fig. 14). HPLC analysis combined with a luminescence assay using the biosensor B_D2HG_-1 confirmed that WbpB also catalyzes the reduction of 2-KG to produce D-2-HG (Fig. 6c,d). As expected, O-antigen polymers of LPS were completely absent when *wbpB* was disrupted in *P. aeruginosa* PAO1 (Δ*wbpB*) (Fig. 6e and Supplementary Fig. 15). Interestingly, these components also clearly decreased in *P. aeruginosa* PAO1 (Δ*PA0317*), indicating that D-2-HG catabolism is also involved in LPS synthesis. The reduction in LPS synthesis in *P. aeruginosa* PAO1 (Δ*PA0317*) might be caused by the accumulated D-2-HG inhibiting WbpB activity (Fig. 6f). In addition to the biosynthesis of L-serine, D-2-HG metabolism is also involved in the generation of LPS in *P. aeruginosa* PAO1 (Fig. 6g).

## Discussion

D-2-HG is able to inhibit the activities of various 2-KG-dependent dioxygenases, transhydrogenase, and transaminase, and influence metabolic homeostasis, cell differentiation, and transaminase, and influence metabolic homeostasis, cell differentiation, and epigenetics^7, 35–38^. However, the mechanism by which organisms respond to D-2-HG and regulate its catabolism has not yet been elucidated. To address this, we took advantage of comparative genomics to identify the D-2-HG responding regulator DhdR in *A. denitrificans* NBRC 15125. We demonstrate that DhdR plays a critical role in controlling the expression of two co-transcribed genes, *dhdR* and *d2hgdh*. EMSA and DNase I footprinting reveal that DhdR directly interacts with a 13 bp inverted repeat in the *dhdR-d2hgdh* promoter region. D-2-HG acts as an antagonist and may prevent the binding of DhdR to the promoter. The dissociation between DhdR and the promoter region induced by D-2-HG is essential to initiate the transcription of *d2hgdh* for D-2-HG catabolism (Fig. 3a).

Besides an oncometabolite, D-2-HG is also an intermediate of various metabolic processes. For example, we previously characterized D-2-HG metabolism in L-serine biosynthesis in *P. stutzeri* A1501. D-2-HG is endogenously produced by SerA to support the thermodynamically unfavorable D-3-PG dehydrogenation in L-serine biosynthesis. D2HGDH then converts D-2-HG back to 2-KG to prevent the toxic effects associated with D-2-HG. Interestingly, SerA in *A. denitrificans* NBRC 15125 also possesses D-2-HG producing activity, but there is no endogenous D-2-HG generation during the growth of this strain. Further investigation confirmed that *A. denitrificans* NBRC 15125 (Δ*serA*) did not exhibit L-serine auxotrophy. L-Serine biosynthesis initiated by SerA is a metabolic process essential for the survival of most organisms. Thus, there might be an unidentified pathway for L-serine biosynthesis in *A. denitrificans* NBRC 15125, that should be further investigated.

Elevated D-2-HG levels are considered to be a biomarker of tumor-associated IDH1/2 mutations and D-2-HGA^1, 2^. Conventional methods to detect D-2-HG are GC-MS/MS and LS-MS/MS, which are time-consuming, laborious, and costly. Recently, enzymatically coupled fluorescence assays were developed to measure the concentration of D-2-HG^14, 39^. NADH produced during the dehydrogenation of D-2-HG by SerA or NAD-dependent D-2-HG dehydrogenase can be oxidized by the diaphorase to produce fluorescent resorufin. The LODs of the enzymatic assays using SerA and NAD-dependent D-2-HG dehydrogenase were 4 μM and 0.13 μM, respectively. In this study, DhdR from *A. denitrificans* NBRC 15125, a regulator that specifically responds to D-2-HG, was used as the biorecognition element to develop a D-2-HG biosensor based on the AlphaScreen technology. An optimized biosensor, B_D2HG_-1, with high sensitivity (LOD = 0.08 μM) and accuracy (Table S4) was generated after introducing 13 point mutations in the DBS. The concentration of D-2-HG spiked into serum and urine, as well as the amount D-2-HG accumulated in different cell lines can be readily determined using B_D2HG_-1 and the results were highly concordant with those of LC-MS/MS. Based on the sensitivity of B_D2HG_-1 and D-2-HG levels in patients, only 1 μL of serum or urine, with no complicated sample pretreatment, was required for clinical diagnosis of D-2-HG-related diseases.

Biosensors are powerful diagnostic devices that combine a biorecognition element with specificity for a given analyte and a transducer element that creates a measurable signal for analysis. In addition to AlphaScreen technology, other strategies such as aTF-NAST (aTF-based nicked DNA-template-assisted signal transduction)^23^, CaT-SMelor (CRISPR-Cas12a and aTF mediated small molecule detector)^24^, and quantum-dot Förster resonance energy transfer (FRET)^25^, can also be coupled with DhdR to construct D-2-HG biosensors. Further, genetically encoded fluorescent probes based on DhdR might also be developed to detect D-2-HG in *vivo*. Beyond reducing 2-KG to D-2-HG, in many organisms, SerA can also reduce oxalacetate to D-malate^13^. Given the importance of L-serine in metabolism, D2HGDH in these organisms has evolved to be a versatile enzyme capable of dehydrogenating both D-malate and D-2-HG to eliminate these two metabolites^40^. Interestingly, D2HGDH in *A. denitrificans* NBRC 15125 has no D-malate dehydrogenation activity and only participates in extracellular D-2-HG catabolism. This enzyme with high specificity toward D-2-HG may also be used as a promising biorecognition element for the development of enzymatic or electrochemical D-2-HG biosensors.

In addition to quantifying D-2-HG in serum and urine, the biosensor B_D2HG_-1 was also used to study bacterial D-2-HG metabolism and revealed that D-2-HG metabolism is involved in L-serine biosynthesis and generation of the O-antigen of LPS in *P. aeruginosa* PAO1. LPS is an important virulence factor of *P. aeruginosa*^34^. Mutant *P. aeruginosa* that are deficient in O-antigen biosynthesis exhibited decreased virulence and was sensitive to serum^41^. Antibiotic resistance in *P. aeruginosa* is a serious threat to human health^42^. AGI-5198 and GSK321 target mutant IDHs and have been developed to treat D-2-HG related cancers^43, 44^. Inhibitors targeting the D-2-HG producing enzyme WbpB, which was identified in this work, might also be developed as narrow-spectrum antibiotics against *P. aeruginosa*. Inactivation of SerA in *P. aeruginosa* significantly inhibits ExoS secretion and penetration through Caco-2 cell monolayers^45^, suggesting that L-serine biosynthesis might also be associated with the pathogenicity of *P. aeruginosa*. Allosteric inhibitors of human SerA have been successfully used to inhibit L-serine biosynthesis in cancer cells and reduce tumor growth^46^. These chemicals might also be effective inhibitors of SerA in *P. aeruginosa* and could potentially be applied as therapies to treat *P. aeruginosa* infection, which is worthy of attempts.

In summary, we uncovered the regulatory mechanism of D-2-HG catabolism in *A. denitrificans* NBRC 15125 and developed a sensitive biosensor to detect D-2-HG in human serum, urine, and cell culture medium. DhdR is the first identified D-2-HG responding regulator which negatively regulates D2HGDH expression and D-2-HG catabolism. Using the biosensor B_D2HG_-1 based on DhdR and AlphaScreen, we also identified the role of D-2-HG metabolism in LPS biosynthesis in *P. aeruginosa* PAO1. The D-2-HG biosensor is thus a valuable experimental tool for revealing other unidentified D-2-HG-related metabolic processes. From a drug-discovery perspective, the fluorometric biosensor with sensitive quantitative capacity also has the potential to be utilized in a high-throughput manner. We hope that the D-2-HG biosensor, like other metabolite biosensors^47–49^, can be used in screening compound, mutation, or siRNA libraries to identify new targets that reduce D-2-HG accumulation for therapeutic purposes.

## Methods

Methods, including statements of data availability and any associated accession codes and references, are available in the online version of the paper.

## Acknowledgments

This work was supported by the grants of National Key R&D Program of China (2019YFA0904800, 2018YFA0901200, and 2019YFA0904900), the National Natural Science Foundation of China (31800025, 31970055), Shandong Provincial Funds for Distinguished Young Scientists (JQ 201806), and Qilu Young Scholar of Shandong University. The funders had no role in study design, data collection and interpretation, or the decision to submit the work for publication. We also thank Zhifeng Li, Jingyao Qu, and Jing Zhu from SKLMT (State Key Laboratory of Microbial Technology, Shandong University) for assistance in isothermal titration calorimetry (ITC) and mass spectrographic analysis.

## Author contributions

C.G., C.M., and P.X. designed the research. D.X., W.Z., X.G., Y.L., C.H., S.G., Z.K., and X.X. performed the research. D.X. and C.G. analyzed the data. D.X., W.Z., C.G., C.M., and P.X. wrote the paper.

## Additional information

Any supplementary information and source data are available in the online version of the paper. Reprints and permissions information is available online at http://www.nature.com/reprints/index.html. Correspondence and requests for materials should be addressed to C.G.

## Online Methods

### Bioinformatics analysis

The distribution of D2HGDH in bacteria were checked by using BLAST as described previously^7^. Individual genomes showing query coverages more than 90%, E values lower than e^−30^, and maximum identity levels higher than 30% with the query protein (D2HGDH in *P. stutzeri* A1501) were selected for further analysis of the upstream and downstream genes of *d2hgdh*.

### Reagents

D-2-Hydroxyglutarate (D-2-HG), L-2-hydroxyglutarate (L-2-HG), D-lactate, L-lactate, D-malate, L-malate, D-tartrate, glutarate, 2-ketoglutarate (2-KG), D-glycerate, L-glycerate, succinate, pyruvate, oxaloacetate, *cis*-aconitate, citrate, isocitrate, dichlorophenol-indophenol (DCPIP), 3-(4,5-dimethylthiazol-2-yl)-2,5-diphenyltetrazolium bromide (MTT), phenazine methosulfate (PMS), NADH, NAD^+^, resazurin, diaphorase, bovine serum albumin (BSA), GSK864, and AGI-6780 used in this study were purchased from Sigma-Aldrich (USA). AlphaScreen donor and acceptor beads were purchased form PerkinElmer (USA). Fumarate and human serum were purchased from Beijing Solarbio Science & Technology Co., Ltd (China). Urine samples were collected from healthy volunteers. D,L-2-Hydroxyglutarate disodium salt (2,3,3-D3) was purchased from Cambridge Isotope Laboratories, Inc. (USA). All other chemicals were of analytical reagent grade.

### Bacteria and culture conditions

Bacterial strains used in this study are listed in Supplementary Table 5. *A. denitrificans* NBRC 15125 and its derivatives were cultured at 30°C and 200 rpm. *E. coli*, *P. aeruginosa* PAO1, and their derivatives were cultured at 37 °C and 180 rpm. Luria-Bertani (LB) medium or minimal medium^50^ containing different carbon sources were used to culture bacterial strains. If necessary, antibiotics were added at appropriate concentrations.

### Mammalian cell culture and treatment by IDH mutant inhibitors

The laboratory-preserved HEK293FT cells were cultured in high-glucose Dulbecco’s modified Eagle’s medium (MACGENE, China) with 10% FBS (Biological Industries (BioInd)) and 1% penicillin/streptomycin (MACGENE, China). HT1080 cells, obtained from Procell Life Science & Technology Co., Ltd, were cultured in Minimum Essential Medium (MEM) medium supplemented with 10% (v/v) fetal bovine serum (FBS) (ThermoFisher, USA) and 1% (v/v) penicillin/streptomycin (MACGENE, China).

The IDH1/R132H-encoding gene was synthesized by Sangon Biotech (China) Co., Ltd, and cloned into the lentiviral vector pLenti PGK GFP Puro (w509-5), replacing the EGFP-encoding gene. The constructed vector was named as PGK-IDH1/R132H-Puro. HEK293FT packaging cells were co-transfected with PGK-IDH1/R132H-Puro, pMD2.G, and psPAX2 by Lipofectamine 3000 (Invitrogen, USA). After 72 h transfection, the supernatant was harvested and filtered through a 0.45 μm filter, and then stored at −80 °C until use. For stable overexpression of IDH1/R132H, HEK293FT cells were infected by above viral supernatant, cultured for further 2 days, and then selected several rounds with 4-6 μg mL^−1^ puromycin. The obtained cells were called as HEK293FT-IDH1/R132H.

GSK864 and AGI-6780 were dissolved in DMSO. HT1080 (3 × 10^5^), HEK293FT (7 × 10^5^), and HEK293FT-IDH1/R132H (7 × 10^5^) cells were seeded into 6-well plates with 2 mL medium per well, respectively. The medium was changed with fresh medium containing 0.5 μM GSK864 or AGI-6780 the next day. After 48 h, the medium was centrifuged at 13,000 × *g* for 5 min, and the obtained supernatant was stored at −80 °C until detection.

### Construction of *A. denitrificans* NBRC 15125 and *P. aeruginosa* PAO1 mutants

The plasmids and primers used in this study are listed in Supplementary Table 5 and Table 6, respectively. To construct the *A. denitrificans* NBRC 15125 (Δ*d2hgdh*), the homologous arms upstream and downstream of the *d2hgdh* gene were amplified with primers *d2hgdh*-uf/*d2hgdh*-ur and *d2hgdh*-df/*d2hgdh*-dr, respectively. The upstream and downstream fragments were fused together by recombinant PCR with primers *d2hgdh*-uf and *d2hgdh*-dr. The generated fusion fragments were digested with *Eco*RI and *Bam*HI and cloned into a suicide plasmid pK18*mobsacB*^51^ cut with the same restriction enzymes. The generated plasmid, pK18*mobsacB*-*d2hgdh’*, was transferred into *A. denitrificans* NBRC 15125 by the tri-parental mating method in the presence of the helper strain *E. coli* HB101 carrying plasmid pRK2013^52, 53^. The single-crossover mutants with integration of the plasmid pK18*mobsacB*-*d2hgdh’* into the chromosome were screened from M9 medium plates containing 20 g L^−1^ trisodium citrate as the sole carbon source and 40 μg mL^−1^ tetracycline. The double-crossover cells were selected on LB plates containing 15% (w/v) sucrose. The *serA* and *dhdR* mutants of *A. denitrificans* NBRC 15125 were constructed by using the same procedure.

*P. aeruginosa* PAO1 (Δ*wbpB*) was constructed based on RedγBAS recombineering system^54^. A DNA fragment containing gentamicin resistance gene flanked by lox71/lox66 sites and 75 bp homologous arms of *wbpB* were amplified with primers *wbpB*-genta-loxM-F/*wbpB*-genta-loxM-R. The PCR products were transferred into the *P. aeruginosa* PAO1 cells, which harbored plasmid pBBR1-Rha-redγBAS-kan and were induced by L-rhamnose. After gentamicin selection (30 μg mL^−1^), the obtained gentamycin-resistant strain was further transformed by the plasmid pCM157. Then, the right colonies were screened from LB plates containing 60 μg mL^−1^ tetracycline, and treated by isopropyl β-D-thiogalactoside (IPTG) induction to eliminate the gentamycin resistance, resulting in the *wbpB* mutant strain *P. aeruginosa* PAO1 (Δ*wbpB*). The *serA* mutant was constructed by using the same procedure. All plasmids and fragments were transferred by electroporation. All the constructed mutants were confirmed by PCR and sequencing.

### Expression, purification and characterization of recombinant D2HGDH, SerA, DhdR, and WbpB

The D2HGDH-encoding gene *d2hgdh* was amplified with primers *d2hgdh*-F/*d2hgdh*-R, which contained *Sac*I and *Hin*dIII restriction enzyme sites, respectively. The PCR product was digested with *Sac*I and *Hin*dIII, and then ligated into the expression vector pETDuet-1. The generated plasmid pETDuet-*d2hgdh* was introduced into *E. coli* BL21(DE3) for D2HGDH expression. To overexpress SerA, DhdR, and WbpB, the corresponding strains were generated by using the same procedure.

The constructed *E. coli* BL21(DE3) strains were grown in LB medium at 37 °C to an OD_600nm_ of 0.6 and induced with 1 mM IPTG at 16 °C for 12 h. The cells were harvested by centrifugation at 6,000 rpm for 10 min at 4 °C and washed twice with buffer A (20 mM sodium phosphate, 20 mM imidazole, and 500 mM NaCl, pH 7.4). The cell pellets were then suspended in the buffer A containing 1 mM phenylmethanesulfonyl fluoride (PMSF) and 10% (v/v) glycerol, lysed by sonication, and centrifuged at 12,000 rpm for 50 min at 4 °C. The supernatant was loaded onto a 5 mL HisTrap HP column (GE Healthcare, USA) pre-equilibrated with buffer A, and then eluted with a gradient ratio of buffer B (20 mM sodium phosphate, 500 mM imidazole, and 500 mM NaCl, pH 7.4) at a flow rate of 5 mL min^−1^. The purified proteins were analyzed by 13% sodium dodecyl sulfate-polyacrylamide gel electrophoresis (SDS-PAGE). Protein concentrations were measured by using the Bradford protein assay kit (Sangon, China).

To determine the native molecular weight of different proteins, size exclusion chromatography was performed using a Superdex 200 10/300 GL column (GE Healthcare, USA). The column was equilibrated with buffer containing 50 mM sodium phosphate and 150 mM NaCl (pH 7.2) at a flow rate of 0.5 mL min^−1^. Thyroglobulin (669 kDa), ferritin (440 kDa), aldolase (158 kDa), conalbumin (75 kDa), ovalbumin (44 kDa), and RNase A (13.7 kDa) were used as standard proteins.

### Enzymatic assays of D2HGDH and SerA

*A. denitrificans* NBRC 15125 were cultured to mid-log stage in minimal medium containing different compounds as carbon sources. Cells were harvested, washed twice, resuspended to an OD_600nm_ of 20 in 50 mM Tris-HCl buffer supplemented with 1 mM PMSF and 10% (v/v) glycerol, then lysed by sonication on ice. The homogenate was centrifuged at 13,000 rpm for 15 min at 4 °C, and the supernatants (the crude cell extracts) were used for D2HGDH activity measurement.

The activity of D2HGDH was assayed at 30 °C by determining the reduction of DCPIP spectrophotometrically at 600 nm in 800 μL reaction mixture containing 50 mM Tris-HCl buffer (pH 7.4), 0.2 mM PMS, 0.05 mM DCPIP, and variable concentrations of substrate. One unit (U) of D2HGDH activity was defined as the amount of enzyme that reduced 1 μmol of DCPIP per minute. The activity of SerA was assayed at 30 °C by measuring the oxidation of NADH spectrophotometrically at 340 nm in 800 μL reaction mixture containing 50 mM Tris-HCl buffer (pH 7.4), 0.2 mM NADH and variable concentrations of 2-KG. One unit (U) of SerA activity was defined as the amount of enzyme that oxidized 1 μmol of NADH per minute. The kinetic constants were calculated by using the double-reciprocal plot method. All the assays were performed using a UV/visible spectrophotometer (Ultrospec 2100 pro, Amersham Biosciences, USA).

### Analysis of the regulation mechanism of DhdR

#### RNA preparation and RT-PCR

Total RNA was extracted from *A. denitrificans* NBRC 15125 cells grown in minimal medium containing D-2-HG as the sole carbon source using the Easypure RNA Kit (Transgen, China). RNA integrity was checked by 1.5% agarose gel electrophoresis. The elimination of genomic DNA contamination and the synthesis of total cDNA was conducted by using HiScript II Q RT SuperMix for qPCR (+gDNA wiper) Kit (Vazyme, China) according to the instructions of manufacturer. Reverse transcription-PCR (RT-PCR) was performed using the total cDNA as template and *d*-*d*-RTF/*d*-*d*-RTR as primers.

#### Determination of the transcriptional start site

The transcription start site of *dhdR*-*d2hgdh* operon was identified by a 5’ rapid amplification of cDNA ends (RACE) system (Invitrogen, China). The first-strand cDNA was synthesized from total RNA with primer GSP1, tailed with terminal deoxynucleotidyl transferase and dCTP, and then amplified by PCR with the abridged anchor primer (APP) and GSP2. A nested PCR was carried out using the PCR product as a template with APP and GSP3. The PCR product was cloned into pMD18-T vector (TaKaRa, China) for sequencing.

#### Electrophoretic mobility shift assay (EMSA)

The DNA fragment F1 were generated by PCR from *A. denitrificans* NBRC 15125 genomic DNA using primers F1-F/F1-R. 10 nM DNA was incubated with purified DhdR (0–100 nM) at 30 °C for 30 min in 20 μL EMSA binding buffer [10 mM Tris-HCl, pH 7.4, 50 mM KCl, 0.5 mM ethylenediaminetetraacetic acid (EDTA), 10% (v/v) glycerol, and 1 mM dithiothreitol (DTT)]. The mixtures were then analyzed by electrophoresis with 6% native polyacrylamide at 4 °C and 170 V (constant voltage) for 45 min. The gel was stained with SYBR green I (TaKaRa, China) for 30 min and photographed by G:BOX F3 gel doc system (Syngene, USA). For the effector analysis of DhdR, purified DhdR was incubated with D-2-HG, L-2-HG, D-malate, D-lactate, glutarate, 2-KG, or pyruvate at 30 °C for 15 min. The DNA fragment (10 nM) was subsequently added into the mixtures and incubated for another 30 min before electrophoresis.

#### DNase I footprinting assay

DNase I footprinting assay was conducted using 6-carboxyfluorescein (FAM) labeled probe and purified DhdR. The fragment F1 was amplified and inserted into pEASY-Blunt Simple Cloning Vector (TransGen, China) to generate pEASY-Blunt-F1. The FAM-labeled probe was amplified with primers M13F-FAM and M13R and plasmid pEASY-Blunt-F1 as template. The PCR product was purified after agarose gel electrophoresis, and quantified by NanoDrop 2000C (Thermo Scientific, USA). For each assay, 400 ng FAM-labeled probe was incubated with 1 μg DhdR in 40 μL EMSA binding buffer at 30 °C for 30 min. Next, 10 μL solution containing about 0.015 U DNase I (Promega, USA) and 100 nmol CaCl2 was added and incubated for 25 °C for 1 min. The reaction was terminated by adding 140 μL stop solution [30 mM EDTA, 200 mM sodium acetate, and 0.15% (w/v) SDS]. Digested samples were subsequently extracted with phenol/chloroform, precipitated with ethanol, dissolved in 30 μL Mini-Q water.

### Development of the biosensors

#### Isothermal titration calorimetry (ITC) experiments

DhdR and the DNA fragments were dissolved in ITC buffer (20 mM sodium phosphate and 300 mM NaCl, pH 8.0). All the DNA fragments were 27 bp in length, obtained by annealing of two reverse complemented primers listed in Supplementary Table 6. Calorimetric assays were conducted on a PEAQ-ITC (MicroCal, USA) at 25 °C with a stir speed of 750 rpm. The reaction cell was filled with 15 μM DhdR solution, and the titration was performed with an initial 0.4 μL injection of 100 μM DNA fragment, followed by 18 injections of 2 μL by 150 s intervals. The control experiment was performed by titrating the corresponding DNA fragment into ITC buffer. The net heat of the dilutions was corrected by subtracting the heat of the control point-to-point. The equilibrium dissociation constant (*K*_D_) was obtained based on a single-site binding model in the assistance of MicroCal PEAQ-ITC Analysis Software.

#### Biotinylated DNA preparation

The biotinylated DNA *dhdO* (Bio-*dhdO*) was prepared by two round PCR. First, unlabeled *dhdO* fragment was amplified via recombinant PCR with the primers *dhdO*-F and *dhdO*-R. Then, Bio-*dhdO* fragment were amplified with primers Bio-F and *dhdO*-R and unlabeled *dhdO* as template. The PCR products were purified after agarose gel electrophoresis, and quantified by NanoDrop ND-1000 (Thermo Scientific, USA). The mutants Bio-*dhdO*-1, -2, -8, and -12 were prepared by using the same procedure.

#### AlphaScreen assays

All the AlphaScreen assays were carried out in 384-well white OptiPlates (PerkinElmer, USA) at room temperature. The 25 μL detecting solution contained 5 μL of each components: DhdR, biotinylated DNA fragment, donor beads, acceptor beads, and samples to be measured. DhdR, biotinylated DNA fragment and samples were mixed and incubated for 30 min. Then, 20 μg mL^−1^ acceptor beads were added and incubated in dark for 30 min. Finally, 20 μg mL^−1^ donor beads were added and incubated for 60 min. The luminescence was measured by EnSight Multimode Plate Reader (PerkinElmer, USA). *I*_50_ was calculated with one-site fit LogI50 models of GraphPad Prism 7.0.

#### Buffer optimization

The buffers used in AlphaScreen assays were described as follows. HBS-P buffer contained 10 mM HEPES and 150 mM NaCl; HBS-EP buffer was HBS-P buffer containing additional 3 mM EDTA; AH buffer contained 25 mM HEPES and 100 mM NaCl; PBS-P buffer contained 100 mM phosphate-buffered saline (PBS). All buffers were adjusted to pH 7.4, supplemented with 0.1% (w/v) BSA and 0.005% Tween-20.

### Preparation and analysis of LPS

LPS was extracted as described previously^55^. Briefly, the cells of *P. aeruginosa* PAO1 and its derivatives grown in LB medium were harvested by centrifugation at 15,600 × g for 1 min, washed three times and resuspended to an OD_600nm_ of 0.45 with PBS. Then, the cell suspension (1 mL) was resuspended in 250 μL lysis buffer [1 M Tris, 10% (v/v) glycerol, and 2% (w/v) SDS, pH 6.8] and heated at 100 °C for 15 min. After adding 30 μg proteinase K and incubating at 55 °C for 5 h, the LPS samples were cooled to room temperature, separated by SDS-PAGE with 13% polyacrylamide gels and visualized by a silver-staining method. 30% (v/v) ethanol-10% (v/v) acetic acid containing 7 g L^−1^ periodic acid was used to oxidize LPS in the gel at 22 °C for 20 min. The gel was washed three times with ddH_2_O for 5 min, stained with 1 g L^−1^ AgNO3 for 30 min at 30 °C. Then, the gel was washed with ddH_2_O for 10 s. The color was developed by 30 g L^−1^ NaCO_3_ and 0.02% (v/v) formaldehyde (pre-cooled on ice). After terminating the color development reaction by 10% (v/v) acetic acid and washing with ddH_2_O, the gel was photographed by CanoScan 9000F MarKII (Cano, Japan).

### Analytical methods

#### High performance liquid chromatography (HPLC) analysis

HPLC system LC-20AT (Shimadzu, Japan) was used to identify the catalytic products in enzymatic reaction samples. The samples were deproteinized before HPLC analysis. In brief, the samples were heated at 100 °C for 15 min. The precipitate of denatured protein was removed by centrifugation at 13,000 rpm for 15 min and being filtered through a 0.22 μm filter. The deproteinized supernatant were analyzed using an Aminex HPX-87H column (300 × 7.8 mm, Bio-Rad, USA) and a refractive index detector. 10 mM H_2_SO_4_ was used as the mobile phase with a flow rate of 0.4 mL min^−1^.

#### Liquid chromatography-tandem mass spectrometry (LC-MS/MS) analysis of D-2-HG

LC-MS/MS system was applied to quantify the D-2-HG in various biological samples. 2,3,3-D3 was supplemented into biological samples as internal standard. Urine samples and cell medium samples were deproteinized with the same procedure used in HPLC analysis; human serum samples were mixed with methanol and vortexed for 2 min, then removed the precipitate of denatured protein as described above. Analysis was performed on a HPLC system LC-20A (Shimadzu, Japan) coupled with a Prominence nano-LTQ Orbitrap velos pro ETD mass spectrometer (ThermoFisher, USA) operating in negative ion mode. Mobile phase consisted of a mixture of methanol and 0.1% triethylamine adjusted to pH 4.5 with acetic acid (5:95). The total run time was 15 min, and 10 μL samples were injected into a Chirobiotic R column (250 × 4.6 mm, Supelco Analytical, USA) with a flow rate of 0.5 mL min^−1^. Monitored transitions for D-2-HG and internal standard were 147.0 → 129.0 m/z and 150.0 → 132.0 m/z, respectively.

## Data availability

The data supporting the findings of this study are available within the article and its Supplementary Information files and from the corresponding authors on request.

