## Supplementary information for "A D-2-hydroxyglutarate biosensor based on specific transcriptional regulator DhdR"

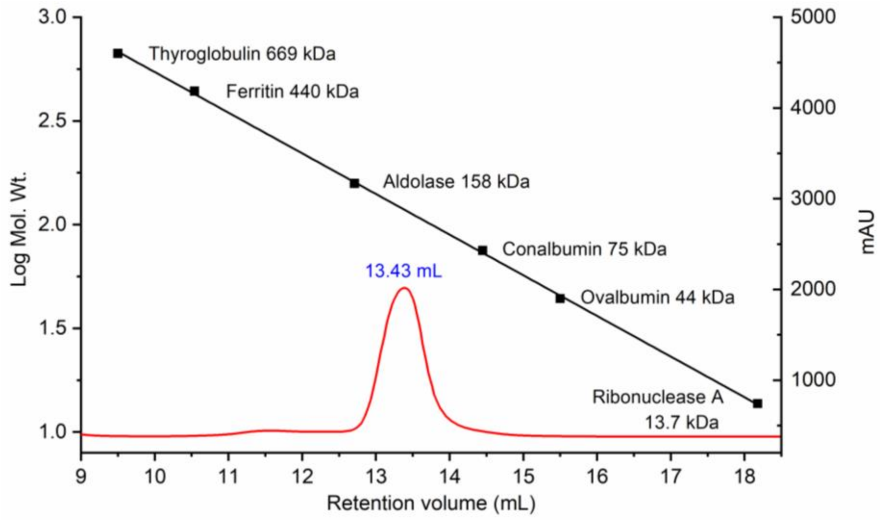


**Supplementary Figure 1. Determination of the molecular weight of D2HGDH by size exclusion chromatography.** With the aid of protein molecular mass standards, a calibration curve (black line) was obtained. The molecular weight of D2HGDH was calculated according to the retention volume (13.43 mL) in the chromatogram (red curve).

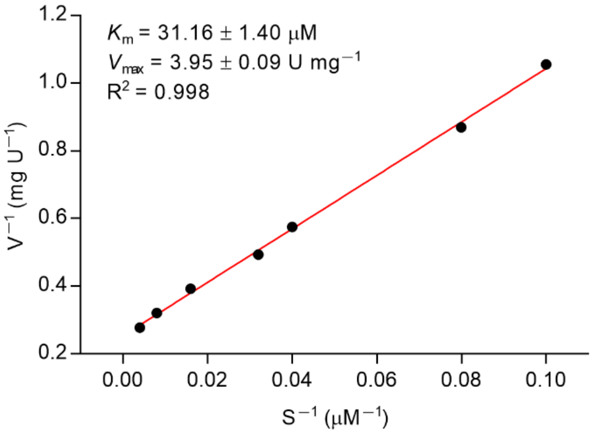


**Supplementary Figure 2. Kinetic parameters of D2HGDH toward D-2-HG.** Data shown are mean ± s.d. ($n$ = 3 independent experiments).

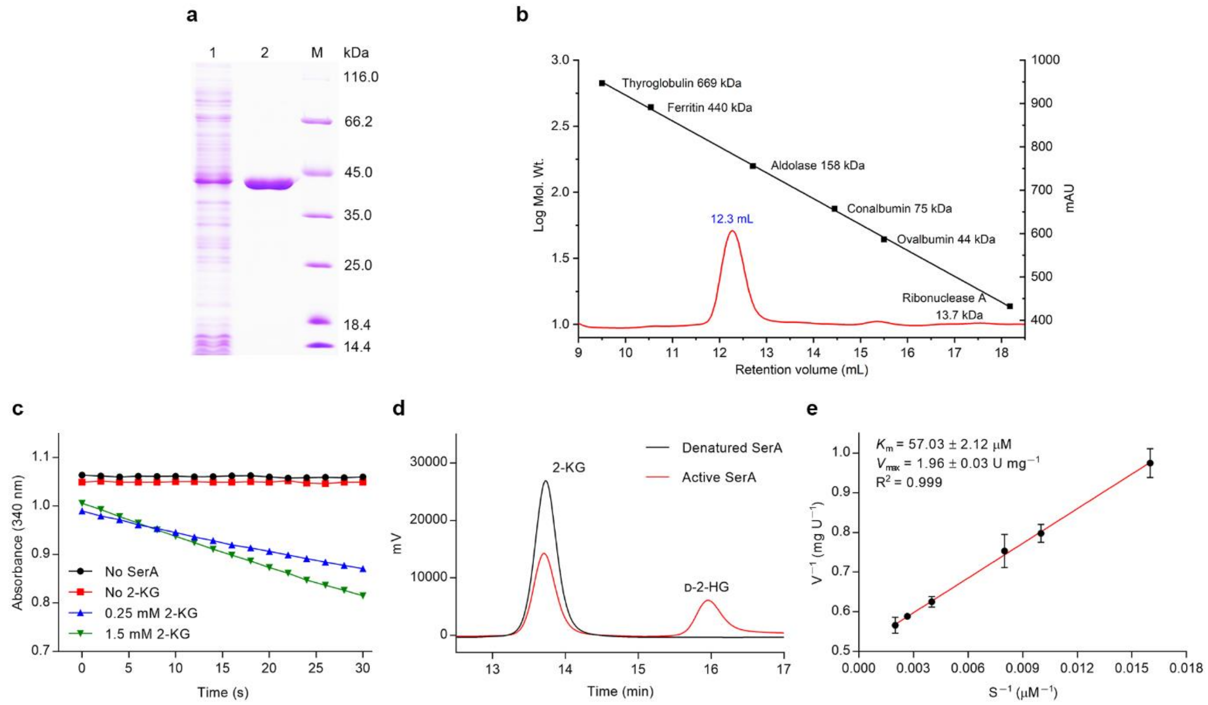


**Supplementary Figure 3. Purification and characterization of SerA. a**, SDS-PAGE analysis of the purified SerA. Lane 1, crude extract of *E. coli* BL21(DE3) harboring pETDuet-*serA*; lane 2, purified SerA using a HisTrap column; lane M, molecular weight markers. **b**, Determination of the molecular weight of SerA by size exclusion chromatography. Black line, standard curve for protein molecular mass standards; red curve, chromatogram of purified SerA. **c**, Activity of SerA toward 2-KG reduction. **d**, HPLC analysis of the product of SerA-catalyzed 2-KG reduction. The reaction mixtures containing 2-KG (50 mM), NADH (40 mM), and active or denatured 2-KG (0.34 mg mL$^{-1}$) in 50 mM Tris-HCl (pH 7.4) were incubated at 37 °C overnight. Black line, the reaction with denatured SerA; red line, the reaction with active SerA. **e**, Kinetic parameters of SerA toward 2-KG. Data shown are mean ± s.d. ($n$ = 3 independent experiments).

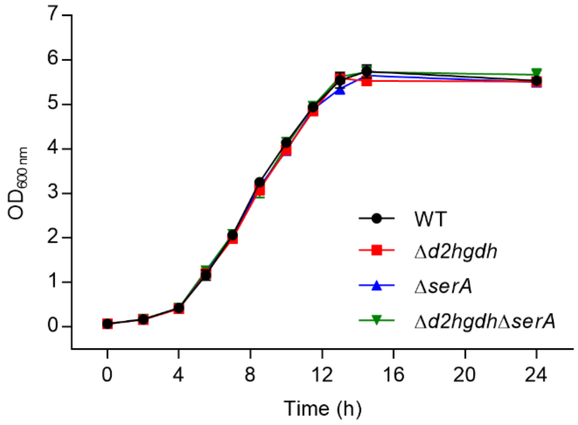


**Supplementary Figure 4. Growth of *A. denitrificans* NBRC 15125 and its derivatives cultured in LB medium.** Data shown are mean ± s.d. (*n* = 3 independent experiments).

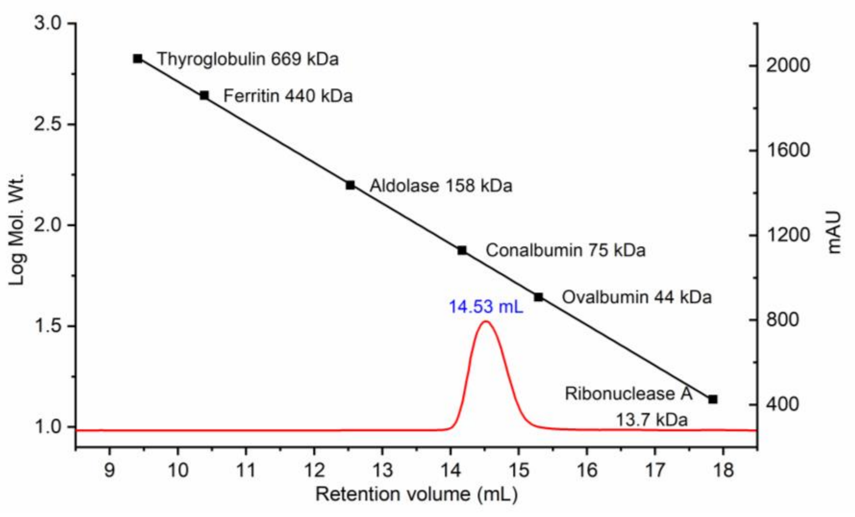


**Supplementary Figure 5. Determination of the molecular weight of DhdR by size exclusion chromatography.** Black line, standard curve for protein molecular mass standards; red curve, chromatogram of purified DhdR.

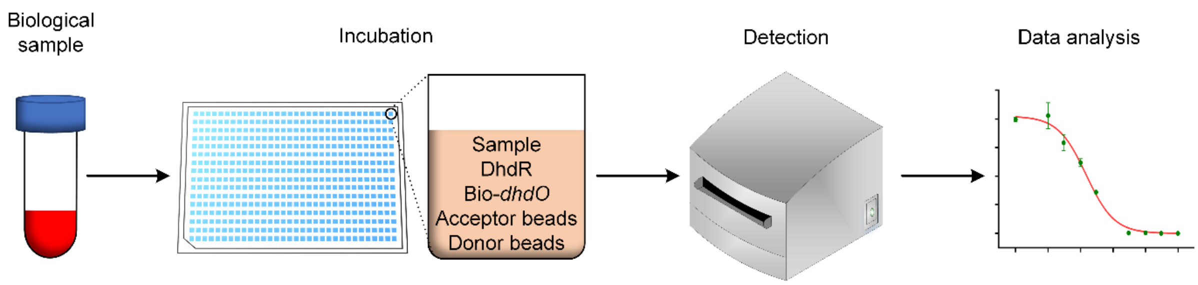


**Supplementary Figure 6. Flow chart for the process of determining D-2-HG in biological sample via the developed biosensor.**

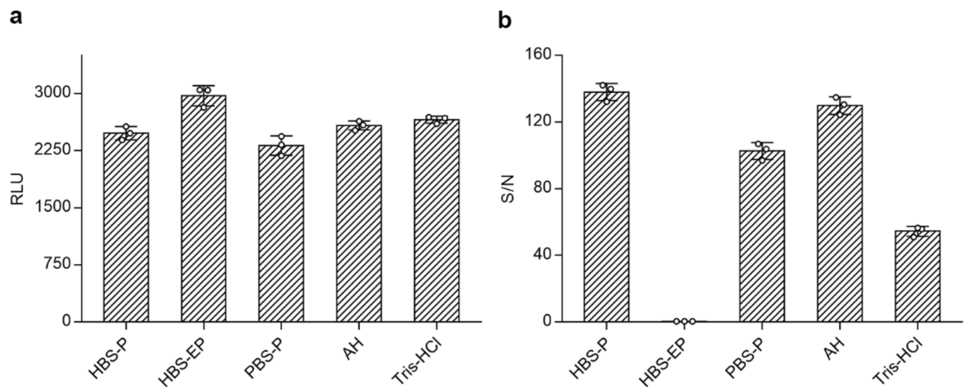


**Supplementary Figure 7. Selection of the optimal buffer for biosensor based on background signal and the ratio of signal to noise (S/N)**. **a**, The background signal of the biosensor in different buffers.The background signals were measured in 25 μL detecting solution containing 5 μL donor beads (20 μg $mL^{-1}$), 5 μL acceptor beads (20 μg $mL^{-1}$) and 15 μL indicated buffer. **b**, The S/N of the biosensor in different buffers. The detecting solution (25 μL) contained 5 μL of each components: DhdR (0.3 nM), Bio-*dhdO* (1 nM), donor beads (20 μg $mL^{-1}$), acceptor beads (20 μg $mL^{-1}$) and corresponding buffer. Data shown are mean ± s.d. ($n$ = 3 independent experiments).

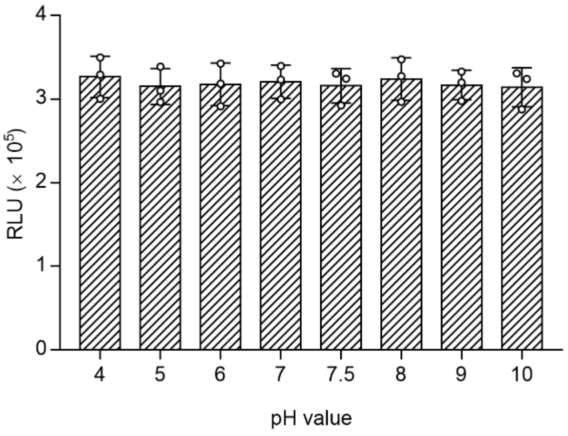


**Supplementary Figure 8. Evaluation of the effect of sample pH on detection.** The detecting solution (25 μL) contained 5 μL of each components: DhdR (0.3 nM), Bio-*dhdO* (1 nM), donor beads (20 μg $mL^{-1}$), acceptor beads (20 μg $mL^{-1}$), and HBS-P buffer with different pH values. Data shown are mean ± s.d. ($n$ = 3 independent experiments).

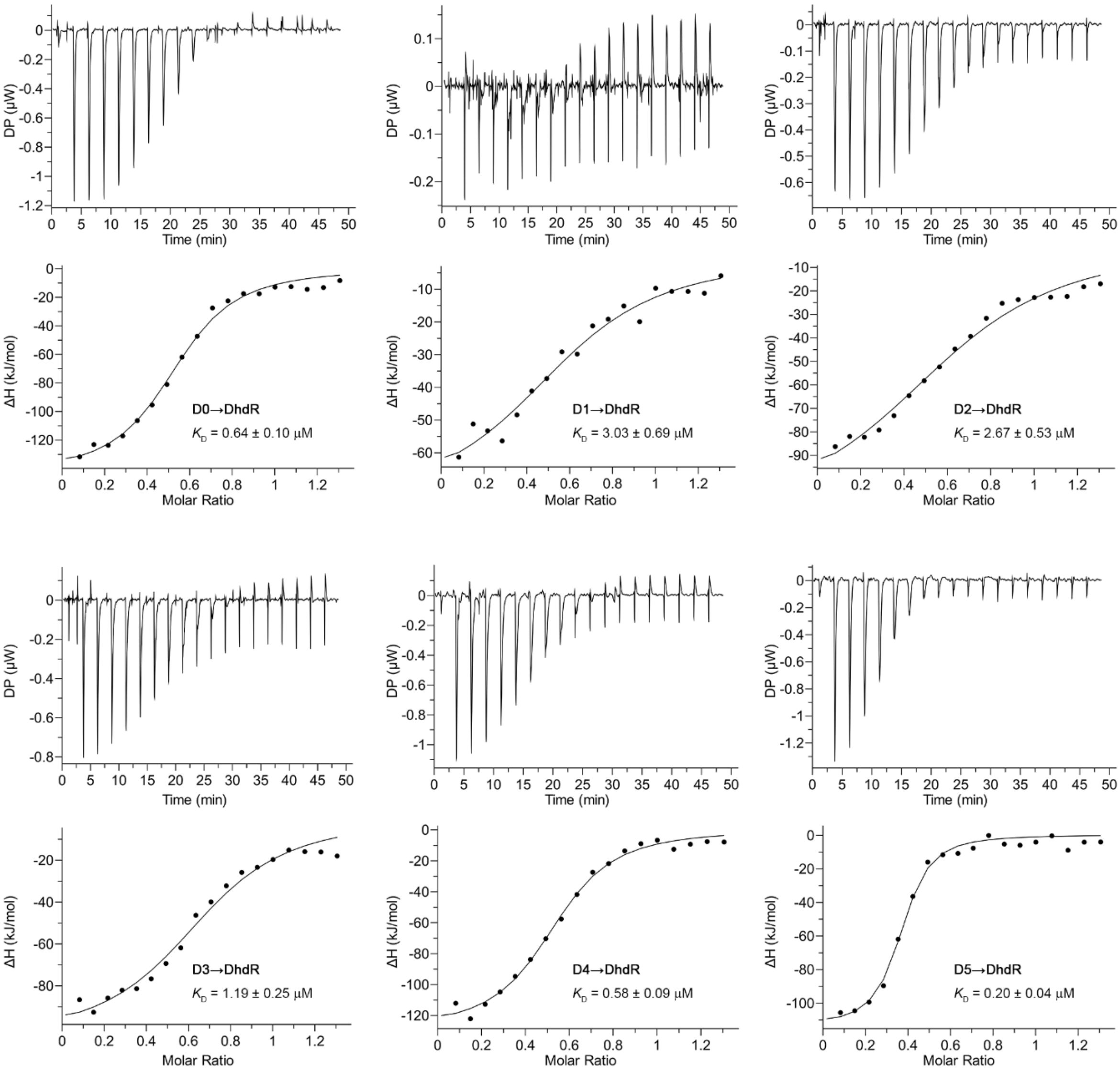


**Supplementary Figure 9-1. Determination of affinities of DhdR with 27-bp DNA fragments containing DBS or DBS mutants by ITC.** 100 μM DNA fragment was titrated to 15 μM DhdR with 19 injections. The control experiment was performed by titrating the corresponding DNA fragment into buffer. No apparent interaction was detected between DhdR and D7, D9, D10, or D11.

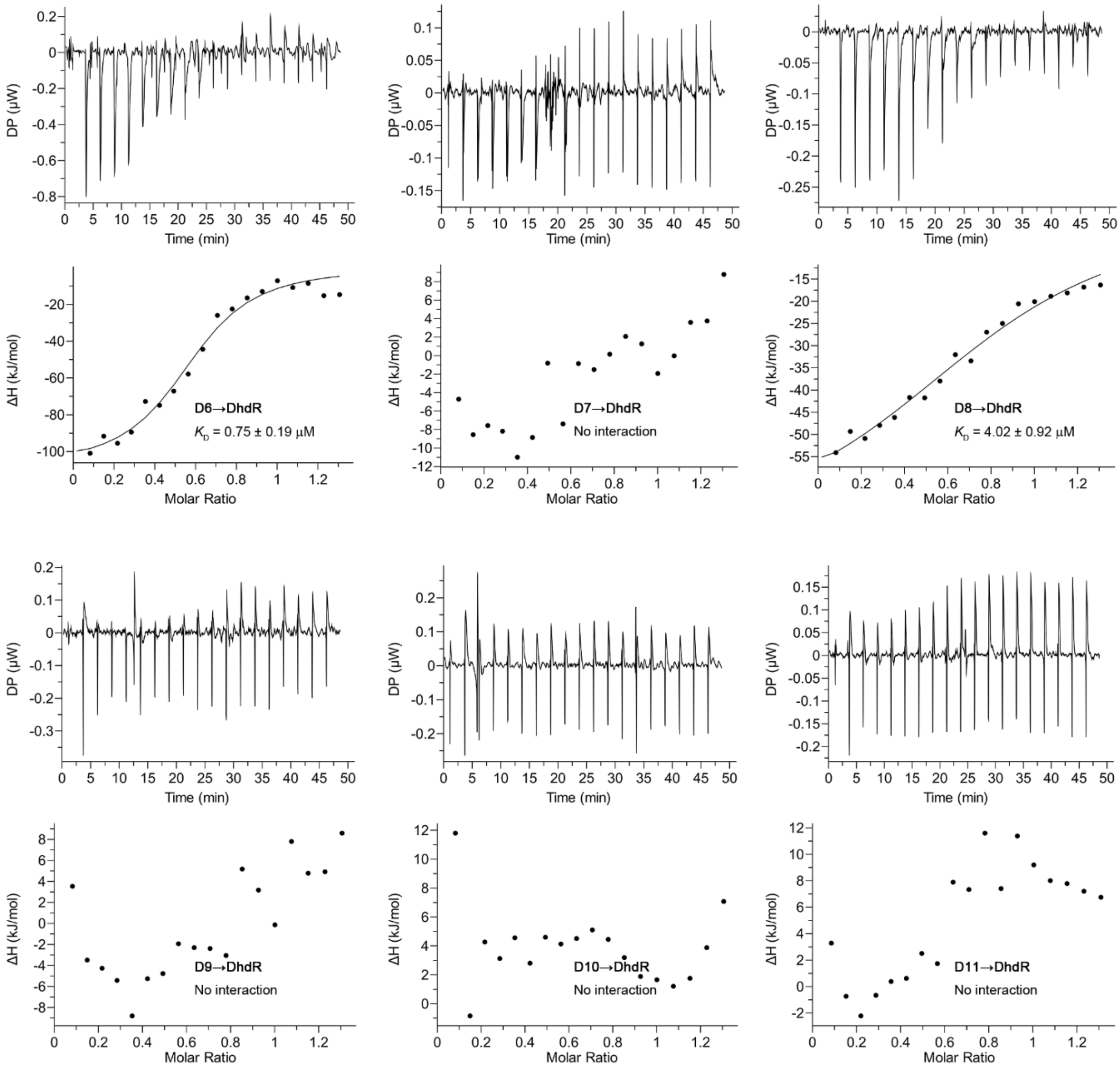


**Supplementary Figure 9-2. Determination of affinities of DhdR for 27-bp DNA fragments containing DBS or DBS mutants by ITC.** 100 μM DNA fragment was titrated to 15 μM DhdR with 19 injections. The control experiment was performed by titrating the corresponding DNA fragment into buffer. No apparent interaction was detected between DhdR and D7, D9, D10, or D11.

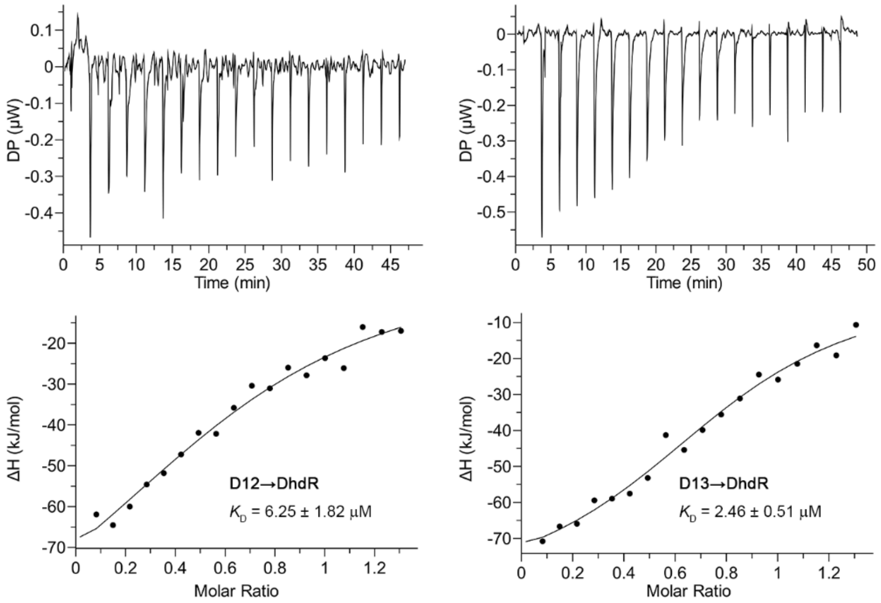


**Supplementary Figure 9-3. Determination of affinities of DhdR for 27-bp DNA fragments containing DBS or DBS mutants by ITC.** 100 μM DNA fragment was titrated to 15 μM DhdR with 19 injections. The control experiment was performed by titrating the corresponding DNA fragment into buffer.

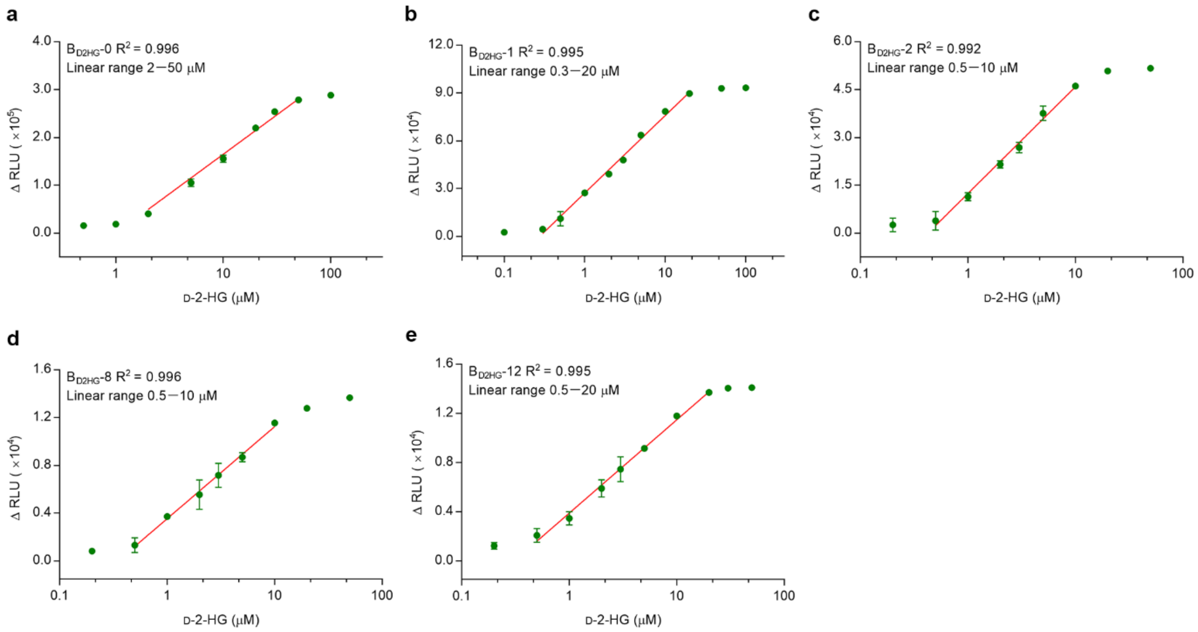


**Supplementary Figure 10. The linear detection ranges of the biosensors. a**, $B_{D2HG}$-0; **b**, $B_{D2HG}$-1; **c**, $B_{D2HG}$-2; **d**, $B_{D2HG}$-8; **e**, $B_{D2HG}$-12. Data shown are mean ± s.d. ($n$ = 3 independent experiments).

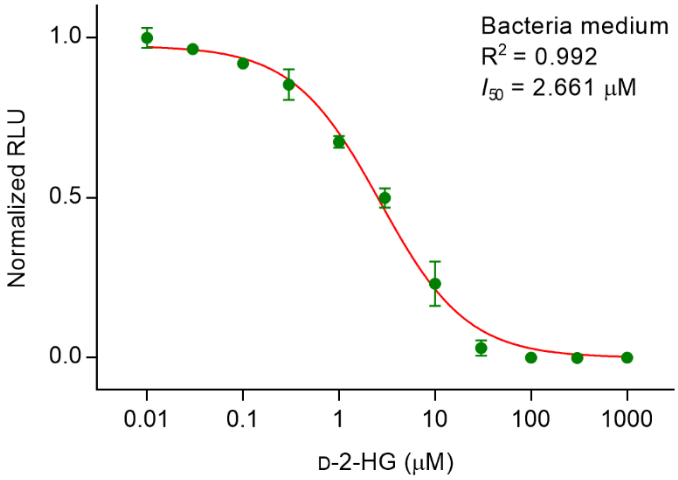


**Supplementary Figure 11. Normalized dose-response curve of $B_{D2HG}$-1 in bacterial minimal medium.** The corresponding luminescence signals are normalized to the maximal value. Data shown are mean ± s.d. ($n$ = 3 independent experiments).

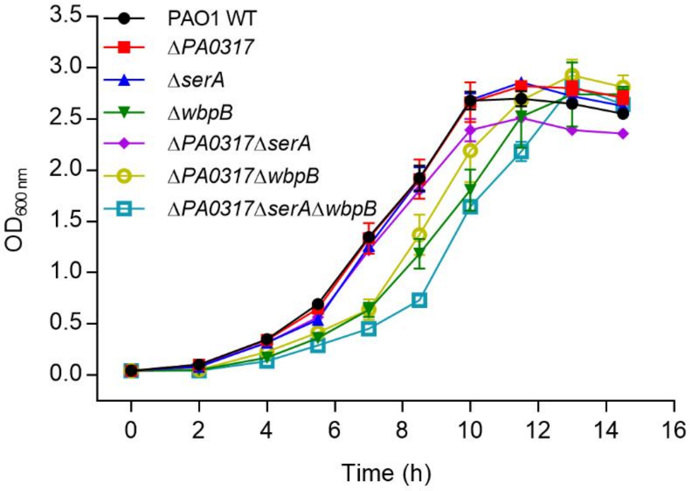


**Supplementary Figure 12. Growth of *P. aeruginosa* PAO1 and its derivatives in minimal medium containing 3 g L$^{-1}$ glucose and 4 mM L-serine.** Data shown are mean ± s.d. ($n$ = 3 independent experiments).

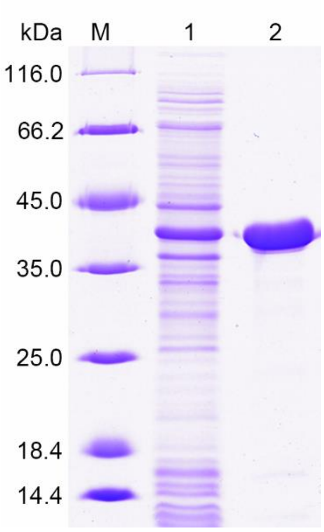


**Supplementary Figure 13. SDS-PAGE analysis of purified WbpB.** Lane M, molecular weight markers; lane 1, crude extract of *E. coli* BL21(DE3) harboring pETDuet-*wbpB*; lane 2, purified WbpB using a HisTrap column.

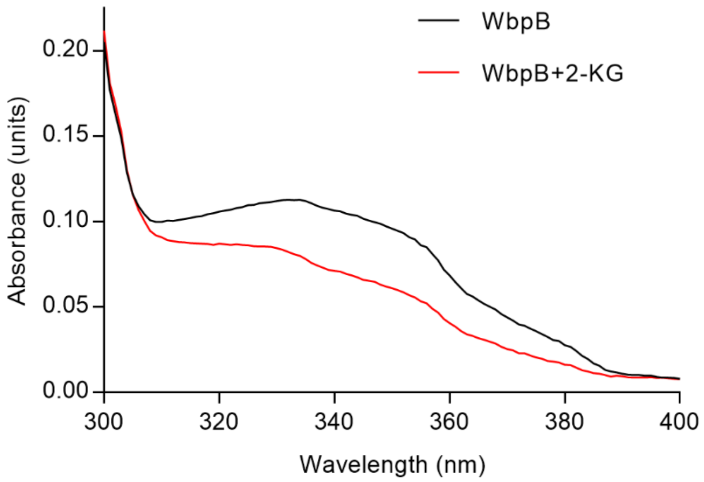


**Supplementary Figure 14. Spectral analysis of WbpB**. The UV-visible absorbance spectra of the purified WbpB was analyzed in 200 μL mixture containing 50 mM Tris-HCl buffer (100 mM NaCl, pH 7.4) and 0.4 mg WbpB before (black line) and after (red line) treated with 1.25 mM 2-KG for 2 h by using an EnSight Multimode Plate Reader (PerkinElmer, USA).

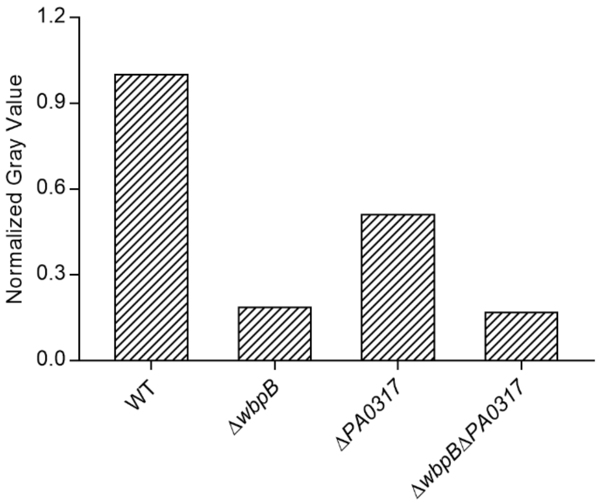


**Supplementary Figure 15. Relative quantification of O-antigen polymers in LPS through gray value analysis of silver-stained SDS-PAGE gel.** Gray value analysis was performed by Image J software.

**Supplementary Table 1** Occurrence of genomic context of *d2hgdh* in bacterial genomes.[a]

| Organism | SerA | GntR | ETF | CadR | LldP | Glycolate oxidase |
|---|---|---|---|---|---|---|
| **Alphaproteobacteria (341)** | **7** | **21** | **4** | **0** | **0** | **0** |
| *Sinorhizobium meliloti* AK83 | – | + | – | – | – | – |
| *Ochrobactrum anthropi* ATCC 49188 | – | + | – | – | – | – |
| *Agrobacterium* sp. H13-3 | – | + | – | – | – | – |
| **Betaproteobacteria (441)** | **2** | **83** | **6** | **0** | **0** | **0** |
| *Bordetella pertussis* 137 | – | + | – | – | – | – |
| *Achromobacter denitrificans* | – | + | – | – | – | – |
| *Achromobacter xylosoxidans* | – | + | – | – | – | – |
| **Deltaproteobacteria (63)** | **7** | **0** | **0** | **0** | **2** | **2** |
| *Corallococcus coralloides* DSM 2259 | + | – | – | – | – | – |
| **Epsilonproteobacteria (159)** | **0** | **0** | **0** | **0** | **0** | **0** |
| **Gammaproteobacteria (351)** | **295** | **1** | **1** | **0** | **0** | **0** |
| *Pseudomonas stutzeri* A1501 | + | – | – | – | – | – |
| *Pseudomonas aeruginosa* PAO1 | + | – | – | – | – | – |
| *Acinetobacter baumannii* | + | – | – | – | – | – |
| **Actinobacteria (276)** | **1** | **4** | **9** | **0** | **1** | **2** |
| *Pseudarthrobacter phenanthrenivorans* Sphe3 | – | + | – | – | + | – |
| **Bacilli (261)** | **0** | **14** | **0** | **70** | **9** | **142** |
| *Bacillus amyloliquefaciens* IT-45 | – | – | – | – | – | + |
| *Bacillus cereus* NJ-W | – | – | – | + | – | + |
| *Bacillus* sp. X1(2014) | – | + | – | – | + | + |
| *Paenibacillus* sp. 32O-W | – | + | – | – | – | + |
| **Clostridia (88)** | **0** | **3** | **35** | **0** | **9** | **3** |
| *Acetohalobium arabaticum* DSM 5501 | – | + | + | – | + | – |
| *Carboxydothermus hydrogenoformans* Z-2901 | – | + | – | – | + | + |
| **Acidobacteria (1)** | **0** | **0** | **0** | **0** | **2** | **0** |
| **Bacteroidetes (35)** | **0** | **0** | **0** | **0** | **0** | **0** |
| **Ignavibacteriae (1)** | **0** | **0** | **0** | **0** | **0** | **1** |
| **Aquificae (16)** | **1** | **0** | **0** | **0** | **0** | **0** |
| **Candidate division NC10 (1)** | **0** | **0** | **0** | **0** | **0** | **0** |
| **Chloroflexi (6)** | **0** | **0** | **0** | **0** | **0** | **0** |
| **Cyanobacteria (20)** | **0** | **0** | **0** | **0** | **0** | **0** |
| **Deferribacteres (1)** | **0** | **0** | **0** | **0** | **0** | **0** |
| **Deinococcus-Thermus (19)** | **0** | **0** | **0** | **0** | **0** | **8** |
| **Erysipelotrichia (1)** | **0** | **0** | **0** | **0** | **0** | **0** |
| **Limnochordia (1)** | **0** | **0** | **0** | **0** | **0** | **1** |
| **Negativicutes (7)** | **0** | **0** | **6** | **0** | **0** | **0** |
| **Tissierellia (1)** | **0** | **0** | **1** | **0** | **0** | **0** |
| **Fusobacteriales (12)** | **0** | **0** | **9** | **0** | **1** | **0** |
| **Nitrospirae (1)** | **0** | **0** | **0** | **0** | **0** | **0** |
| **Acidithiobacillia (5)** | **0** | **0** | **0** | **0** | **0** | **0** |
| **Spirochaetes (16)** | **0** | **0** | **4** | **0** | **0** | **0** |
| **Synergistetes (2)** | **0** | **0** | **2** | **0** | **0** | **0** |
| **Tenericutes (2)** | **0** | **0** | **0** | **0** | **0** | **0** |
| **Thermobaculum (1)** | **0** | **0** | **0** | **0** | **0** | **0** |
| **Thermodesulfobacteria (4)** | **0** | **0** | **0** | **0** | **1** | **0** |
| **Thermotogae (6)** | **0** | **0** | **6** | **0** | **0** | **0** |
| **Verrucomicrobia (4)** | **1** | **0** | **0** | **0** | **0** | **0** |
| **Total: 2143** | **314** | **126** | **83** | **70** | **25** | **159** |

[a]Representative species in several taxonomic groups of bacteria are shown as rows and the presence or absence of genes enconding the respective functional protein (columns) is shown by + or −. Numbers for taxonomic group rows indicate the number of species that have a gene ortholog.

**Supplementary Table 2** Determination of the influence of foreign substance on luminescence signal.

| Foreign substance | Change of luminescence signal[a] (%) | Foreign substance | Change of luminescence signal (%) |
|---|---|---|---|
| $K^+$ | 2.94 | Creatinine | 0.15 |
| $NH_4^+$ | 0.97 | Sucrose | 0.49 |
| $Na^+$ | 0.06 | Urea | 1.31 |
| $Ca^{2+}$ | 0.81 | Ascorbic | 1.17 |
| $Mg^{2+}$ | 2.07 | Leucine | 1.99 |
| Methionine | 0.54 | Valine | 1.35 |
| Alanine | 0.51 | Serine | 0.72 |

[a]Data of change of luminescence signal shown are the mean value (*n* = 3 independent experiments).

**Supplementary Table 3** Key parameters of the developed biosensors.

| Parameter | $B_{D2HG}$-0 | $B_{D2HG}$-1 | $B_{D2HG}$-2 | $B_{D2HG}$-8 | $B_{D2HG}$-12 |
|---|---|---|---|---|---|
| $K_D$ (μM)[a] | 0.64 | 3.03 | 2.67 | 4.02 | 6.25 |
| $I_{50}$ (μM)[b] | 9.05 | 1.59 | 2.69 | 1.62 | 1.68 |
| LOD (μM)[c] | 0.80 | 0.08 | 0.47 | 0.24 | 0.29 |
| Detection range (μM) | 2–50 | 0.3–20 | 0.5–10 | 0.5–10 | 0.5–20 |

[a] $K_D$ was determined by ITC and analyzed by using a single-site binding model.

[b] $I_{50}$ was calculated with one-site fit $LogI_{50}$ models of GraphPad Prism 7.0.

[c] LOD was calculated by interpolating the $RLU_{LOD}$ into the dose-reponse curve. $RLU_{LOD}$=average $B'_{max}(3\text{ wells}) - 3 \times SD(3\text{ wells})$.

**Supplementary Table 4** Evaluation of the performance of $B_{D2HG}$-1 in biological samples.

| Condition | Approach | Concentration (μM)[a] | | | Accuracy (%)[b] | | | Precision |
|---|---|---|---|---|---|---|---|---|
| | Standard | 150 | 1500 | 3500 | 150 | 1500 | 3500 | (RSD%)[c] |
| Serum | LC-MS/MS | 142.21 | 1596.44 | 3629.37 | 94.81 | 106.43 | 103.70 | 5.98 |
| | $B_{D2HG}$-1 | 135.42 | 1494.89 | 3737.51 | 106.13 | 104.62 | 101.43 | 2.31 |
| Urine | LC-MS/MS | 159.39 | 1370.13 | 3332.71 | 106.26 | 91.34 | 95.22 | 7.93 |
| | $B_{D2HG}$-1 | 145.41 | 1456.73 | 3577.44 | 107.12 | 103.10 | 98.13 | 4.38 |
| Cell medium | LC-MS/MS | 173.59 | 1777.85 | 4512.84 | 115.73 | 118.52 | 128.94 | 5.75 |
| | $B_{D2HG}$-1 | 150.49 | 1378.03 | 3740.92 | 99.42 | 99.63 | 102.10 | 1.49 |

[a]Concentration was the data detected from biological samples by $B_{D2HG}$-1. The standard concentrations were the spiked concentrations to biological sample.

[b]Accuarcy% = $\frac{\text{Concentraion determined by LC-MS/MS or } B_{D2HG}\text{-1}}{\text{Standard concentration}}$.

[c]Precision% = $\frac{\text{Standard derivation of accuracy}}{\text{Mean value of accuracy}}$.

**Supplementary Table 5** Strains and plasmids used in this study.

| Strain or plasmid | Relevant characteristics[a] |
|---|---|
| ***A. denitrificans*** | |
| NBRC 15125 | Wild-type |
| NBRC 15125 (Δ*d2hgdh*) | NBRC 15125 with a deletion of the *d2hgdh* gene |
| NBRC 15125 (Δ*serA*) | NBRC 15125 with a deletion of the *serA* gene |
| NBRC 15125 (Δ*d2hgdh*Δ*serA*) | NBRC 15125 with deletions of the *d2hgdh* gene and *serA* gene |
| NBRC 15125 (Δ*dhdR*) | NBRC 15125 with a deletion of the *dhdR* gene |
| ***P. aeruginosa*** | |
| PAO1 | Wild-type |
| PAO1 (Δ*PA0317*) | PAO1 with a deletion of the *PA0317* gene |
| PAO1 (Δ*serA*) | PAO1 with a deletion of the *serA* gene |
| PAO1 (Δ*wbpB*) | PAO1 with a deletion of the *wbpB* gene |
| PAO1 (Δ*PA0317*Δ*serA*) | PAO1 with deletions of the *PA0317* gene and *serA* gene |
| PAO1 (Δ*PA0317*Δ*wbpB*) | PAO1 with deletions of the *PA0317* gene and *wbpB* gene |
| PAO1 (Δ*PA0317*Δ*serA*Δ*wbpB*) | PAO1 with deletions of the *PA0317* gene, *serA* gene and *wbpB* gene |
| ***E. coli*** | |
| DH5α | $F^-$ φ80*lacZ*ΔM15 Δ(*lacZYA*-*argF*)U169 *deoR recA*1 *endA*1 *hsdR*17($r_K^-$, $m_K^+$) *phoA supE*44 $\lambda^-$ *thi*-1 *gyrA*96 *relA*1, used for gene clone |
| BL21(DE3) | $F^-$ *ompT hsdSB*(*rB- mB-*) *gal*(λ *c I* 857 *ind1 Sam*7 *nin*5 *lac*UV5-T7*gene*1) *dcm* (DE3) |
| HB101 | $F^-$ *mcrB mrr hsdS20*($r_B^-$ $m_B^-$) *recA13 supE44 ara14 proA2 lacY1 galK2 xy15* $\lambda^-$ *mtl1 rpsL20*($Sm^r$) *glnV44* $\lambda^-$; triparental mating helper strain |
| BL21-D2HGDH | *E. coli* BL21(DE3) harboring the expression plasmid pETDuet-*d2hgdh* |
| BL21-SerA | *E. coli* BL21(DE3) harboring the expression plasmid pETDuet-*serA* |
| BL21-DhdR | *E. coli* BL21(DE3) harboring the expression plasmid pETDuet-*dhdR* |
| BL21-WbpB | *E. coli* BL21(DE3) harboring the expression plasmid pETDuet-*wbpB* |
| **Plasmid** | |

| | |
|---|---|
| pETDuet-1 | Ap$^r$, vector for protein expression |
| pETDuet-*d2hgdh* | Ap$^r$, pETDuet-1 contained *d2hgdh* gene of *A. denitrificans* NBRC 15125 |
| pETDuet-*serA* | Ap$^r$, pETDuet-1 contained *serA* gene of *A. denitrificans* NBRC 15125 |
| pETDuet-*dhdR* | Ap$^r$, pETDuet-1 contained *dhdR* gene of *A. denitrificans* NBRC 15125 |
| pETDuet-*wbpB* | Ap$^r$, pETDuet-1 contained *wbpB* gene of *P. aeruginosa* PAO1 |
| pRK2013 | Km$^r$, ColE1 ori, *mob*, *tra*$^+$; helper plasmid for conjugation experiments |
| pK18*mobsacB* | Km$^r$, suicide plasmid for gene knockout |
| pK18*mobsacB*-*d2hgdh'* | Km$^r$, partial lengths of *d2hgdh* were inserted into pK18*mobsacB* |
| pK18*mobsacB*-*serA'* | Km$^r$, partial lengths of *serA* were inserted into pK18*mobsacB* |
| pK18*mobsacB*-*dhdR'* | Km$^r$, partial lengths of *dhdR* were inserted into pK18*mobsacB* |
| pMD18-T | Ap$^r$, TA cloning vector |
| pEASY-Blunt cloning vector | Ap$^r$, vector for gene cloning |
| pEASY-Blunt-F1 | Ap$^r$, pEASY-Blunt cloning vector with 81-bp fragment upstream of *dhdR* |
| pBBR1-Rha-redγBAS-kan | Km$^r$, BAS gene and redγ under Rha promoter |
| pR6K-Tps-gentaR-tetR-T7RP | Gm$^r$, PCR templates to amplify lox71-gentaR-lox66 |
| pCM157 | Tc$^r$, pCM62 with *cre* from pJW168; *cre* expression vector |
| pLenti PGK GFP Puro (w509-5) | Ap$^r$, lentiviral expression vector |
| PGK-IDH1R132H-Puro | Ap$^r$, the plasmid carrying IDH1R132H-encoding gene using pLenti PGK GFP Puro (w509-5) as backbone |
| pMD2.G | Ap$^r$, VSV-G envelope expressing plasmid |
| psPAX2 | Ap$^r$, lentiviral packaging vector |

[a]Ap$^r$, ampicillin resistant; Km$^r$, kanamycin resistant; Gm$^r$, gentamicin resistant; Tc$^r$, tetracycline resistant.

**Supplementary Table 6** Primers used in this work.

| Primer | Sequence (5'-3')[a] | Use |
|---|---|---|
| **Overexpression** | | |
| *d2hgdh*-F | ATAT*GAGCTC*GATGAGCGCATCCGACTTTA (*Sac*I) | Amplification of *d2hgdh* in *A. denitrificans* NBRC 15125 (forward) |
| *d2hgdh*-R | TATT*AAGCTT*CTACAGCAGCTTCCCGGGAT (*Hin*dIII) | Amplification of *d2hgdh* in *A. denitrificans* NBRC 15125 (reverse) |
| *serA*-F | TACT*GAATTC*GATGGCACAAATCGTCCTGTT (*Eco*RI) | Amplification of *serA* in *A. denitrificans* NBRC 15125 (forward) |
| *serA*-R | TCGC*AAGCTT*CTACAGCCGGTCGCAG (*Hin*dIII) | Amplification of *serA* in *A. denitrificans* NBRC 15125 (reverse) |
| *dhdR*-F | ATCT*GAGCTC*GATGCTGAGCAAGAGCCTG (*Sac*I) | Amplification of *dhdR* in *A. denitrificans* NBRC 15125 (forward) |
| *dhdR*-R | TATT*AAGCTT*TCATGATGTCTGCCTTGCGG (*Hin*dIII) | Amplification of *dhdR* in *A. denitrificans* NBRC 15125 (reverse) |
| *wbpB*-F | ATAT*GGATCC*GATGAAAAATTTCGC (*Bam*HI) | Amplification of *wbpB* in *P. aeruginosa* PAO1 (forward) |
| *wbpB*-R | ATAT*AAGCTT*TCAACGCGCAAGCGC (*Hin*dIII) | Amplification of *wbpB* in *P. aeruginosa* PAO1 (reverse) |
| **Gene knockout** | | |
| *d2hgdh*-uf | CTATGACATGATTAC*GAATTC*ATGAGCGCATCCGACTTTACG (*Eco*RI) | Amplification of upstream homologous arm of *d2hgdh* (forward) |
| *d2hgdh*-ur | GACCGCATGAACGCCGTGCGTTTGTACCCGGACATCCGCCC | Amplification of upstream homologous arm of *d2hgdh* (reverse) |
| *d2hgdh*-df | GGGCGGATGTCCGGGTACAAACGCACGGCGTTCATGCGGTC | Amplification of downstream homologous arm of *d2hgdh* (forward) |
| *d2hgdh*-dr | TCGCTAACGGATTCA*GGATCC*CTACAGCAGCTTCCCGG (*Bam*HI) | Amplification of downstream homologous arm of *d2hgdh* (reverse) |
| *serA1*-uf | CTATGACATGATTAC*GAATTC*ATGGCACAAATCGTCCTG (*Eco*RI) | Amplification of upstream homologous arm of *serA* (forward) |
| *serA1*-ur | CACCAGCTTCTCGGCGACTTATGGCTTCGCCCAGCACCA | Amplification of upstream homologous arm of *serA* (reverse) |
| *serA1*-df | TGGTGCTGGGCGAAGCCATAAGTCGCCGAGAAGCTGGTG | Amplification of downstream homologous arm of *serA* (forward) |
| *serA1*-dr | TCGCTAACGGATTCA*GGATCC*CTACAGCCGGTCGCAGCG (*Bam*HI) | Amplification of downstream homologous arm of *serA* (reverse) |
| *dhdR*-uf | CTATGACATGATTAC*GAATTC*GGCGGACGCTGCTGCGCGGTCG (*Eco*RI) | Amplification of upstream homologous arm of *dhdR* (forward) |
| *dhdR*-ur | CCTGACGGCGGCTGTTGGTGCGCGCTCACACCGTACTGCTC | Amplification of upstream homologous arm of *dhdR* (reverse) |

| | | |
|---|---|---|
| *dhdR*-df | GAGCAGTACGGTGTGAGCGCGCACCAACAGCCGCCGTCAGG | Amplification of downstream homologous arm of *dhdR* (forward) |
| *dhdR*-dr | TCGCTAACGGATTCA*GGATCC*TGGCCCTTGTACAGGCCGC (*Bam*HI) | Amplification of downstream homologous arm of *dhdR* (reverse) |
| *serA2*-genta-loxM-F | CCGGCCGGACCCGCTTCGCGCAGTTCGCTTTCCAGCCACTTCCTCCGGGACAACAGGTTTACGCAGATGAGCAAGAGCTGAATTACATTCCCAACCG | Amplification of homologous arm of *serA* in *P. aeruginosa* PAO1 and gentamicin resistance gene (forward) |
| *serA2*-genta-loxM-R | AAGGAGGCCGCTGGGCCTCCTTTTCCTTTCCGCCGGACGCTGCCTTAGAACAGCACGCGGCTACGGATGGTACCGCAACTTAAATGTGAAAGTGGGT | Amplification of homologous arm of *serA* in *P. aeruginosa* PAO1 and gentamicin resistance gene (reverse) |
| *serA2*-F | ACAACTCAGGTCGCAACCGGGCA | Verification of deletion of *serA* in *P. aeruginosa* PAO1 (forward) |
| *serA2*-R | ACTTCGGCAAGGGCCAGACC | Verification of deletion of *serA* in *P. aeruginosa* PAO1 (reverse) |
| *wbpB*-genta-loxM-F | GATCACCCATCCCAGCATGTCCATCCGCTCGTGCCAGAAGGCCGGGCGGATCCGCTCATTTCCATAGGACGAACCAGCTGAATTACATTCCCAACCG | Amplification of homologous arm of *wbpB* in *P. aeruginosa* PAO1 and gentamicin resistance gene (forward) |
| *wbpB*-genta-loxM-R | ATGACATGCTGATGAATGATCCTGCGGAAACCTGCTGTAAACAGGTGACCGAGGACGGCCACCTCCTTTTCTACCCAACTTAAATGTGAAAGTGGGT | Amplification of homologous arm of *wbpB* in *P. aeruginosa* PAO1 and gentamicin resistance gene (reverse) |
| *wbpB*-F1 | GACAAGTTTGACTATGAGCTGATC | Verification of the deletion of *wbpB* in *P. aeruginosa* PAO1 (forward) |
| *wbpB*-R1 | AAGCTAGCACAGCCAGAACT | Verification of the deletion of *wbpB* in *P. aeruginosa* PAO1 (reverse) |
| **Cotranscription assay** | | |
| *d*-*d*-RTF | CGCACCAACAGCCGCCGTCA | Amplification of overlapping regions spanning the *dhdR*-*d2hgdh* genes (forward) |
| *d*-*d*-RTR | TGGCCCTTGTACAGGCCGCGCCA | Amplification of overlapping regions spanning the *dhdR*-*d2hgdh* genes (reverse) |
| **RACE-PCR** | | |
| GSP1 | GGCGACGTGGAAGGCGAT | Oligonucleotide for reverse transcription |
| GSP2 | CGCCCTGTTCCGGATGGT | Oligonucleotide for first round PCR of the 5′ RACE |
| GSP3 | CGCCGCCTTCCAGTTCCAT | Oligonucleotide for nested PCR of the 5′ RACE |

| | | |
|---|---|---|
| **EMSA** | | |
| F1-F | GAGTCGCGGCGGCGCGCCGGAT | Amplification of fragment F1 (forward) |
| F1-R | GCGCCGATTATAGGCCTACTT | Amplification of fragment F1 (reverse) |
| **ITC**[b] | | |
| D0-F | AAAGTTATCAGATAACCTGAAAAGTAG | Amplification of fragment D0 (forward) |
| D0-R | CTACTTTTCAGGTTATCTGATAACTTT | Amplification of fragment D0 (reverse) |
| D1-F | AAATTTATCAGATAACCTGAAAAGTAG | Amplification of fragment D1 (forward) |
| D1-R | CTACTTTTCAGGTTATCTGATAAATTT | Amplification of fragment D1 (reverse) |
| D2-F | AAAGGTATCAGATAACCTGAAAAGTAG | Amplification of fragment D2 (forward) |
| D2-R | CTACTTTTCAGGTTATCTGATACCTTT | Amplification of fragment D2 (reverse) |
| D3-F | AAAGTGATCAGATAACCTGAAAAGTAG | Amplification of fragment D3 (forward) |
| D3-R | CTACTTTTCAGGTTATCTGATCACTTT | Amplification of fragment D3 (reverse) |
| D4-F | AAAGTTCTCAGATAACCTGAAAAGTAG | Amplification of fragment D4 (forward) |
| D4-R | CTACTTTTCAGGTTATCTGAGAACTTT | Amplification of fragment D4 (reverse) |
| D5-F | AAAGTTAGCAGATAACCTGAAAAGTAG | Amplification of fragment D5 (forward) |
| D5-R | CTACTTTTCAGGTTATCTGCTAACTTT | Amplification of fragment D5 (reverse) |
| D6-F | AAAGTTATAAGATAACCTGAAAAGTAG | Amplification of fragment D6 (forward) |
| D6-R | CTACTTTTCAGGTTATCTTATAACTTT | Amplification of fragment D6 (reverse) |
| D7-F | AAATGTATCAGATAACCTGAAAAGTAG | Amplification of fragment D7 (forward) |
| D7-R | CTACTTTTCAGGTTATCTGATACATTT | Amplification of fragment D7 (reverse) |
| D8-F | AAATTGATCAGATAACCTGAAAAGTAG | Amplification of fragment D8 (forward) |
| D8-R | CTACTTTTCAGGTTATCTGATCAATTT | Amplification of fragment D8 (reverse) |
| D9-F | AAATTTCTCAGATAACCTGAAAAGTAG | Amplification of fragment D9 (forward) |
| D9-R | CTACTTTTCAGGTTATCTGAGAAATTT | Amplification of fragment D9 (reverse) |
| D10-F | AAATTTAGCAGATAACCTGAAAAGTAG | Amplification of fragment D10 (forward) |
| D10-R | CTACTTTTCAGGTTATCTGCTAAATTT | Amplification of fragment D10 (reverse) |
| D11-F | AAATTTATAAGATAACCTGAAAAGTAG | Amplification of fragment D11 (forward) |
| D11-R | CTACTTTTCAGGTTATCTTATAAATTT | Amplification of fragment D11 (reverse) |

| | | |
|---|---|---|
| D12-F | AAAGGGATCAGATAACCTGAAAAGTAG | Amplification of fragment D12 (forward) |
| D12-R | CTACTTTTCAGGTTATCTGATCCCTTT | Amplification of fragment D12 (reverse) |
| D13-F | AAAGGTCTCAGATAACCTGAAAAGTAG | Amplification of fragment D13 (forward) |
| D13-R | CTACTTTTCAGGTTATCTGAGACCTTT | Amplification of fragment D13 (reverse) |
| **Biotinylated DNA**[c] | | |
| Bio-F | GAGTCGCGGCGGCGCGCCGGAT | Amplification of biotinylated DNA fragment (forward) |
| *dhdO*-F | GAGTCGCGGCGGCGCGCCGGATCCGGGCTGTCATTGTCA | Amplification of fragment *dhdO* (forward) |
| *dhdO*-R | GCGCCGATTATAGGCCTACTTTTCAGGTTATCTGATAACTTTT<br>GACAATGACAGCCCGGAT | Amplification of fragment *dhdO* (reverse) |
| *dhdO*-1-R | GCGCCGATTATAGGCCTACTTTTCAGGTTATCTGATAAATTTT<br>GACAATGACAGCCCGGAT | Amplification of fragment *dhdO*-1 (reverse) |
| *dhdO*-2-R | GCGCCGATTATAGGCCTACTTTTCAGGTTATCTGATACCTTTT<br>GACAATGACAGCCCGGAT | Amplification of fragment *dhdO*-2 (reverse) |
| *dhdO*-8-R | GCGCCGATTATAGGCCTACTTTTCAGGTTATCTGATCAATTTT<br>GACAATGACAGCCCGGAT | Amplification of fragment *dhdO*-8 (reverse) |
| *dhdO*-12-R | GCGCCGATTATAGGCCTACTTTTCAGGTTATCTGATCCCTTTT<br>GACAATGACAGCCCGGAT | Amplification of fragment *dhdO*-12 (reverse) |

[a]Restriction sites are in italic, and the restriction enzymes are indicated in parentheses.

[b]The mutant sites are underlined.

[c]For the amplification of fragment *dhdO*, *dhdO*-1, *dhdO*-2, *dhdO*-8, and *dhdO*-12, primer *dhdO*-F was used as the forward primer.
